# Oncogenic Ras-Src-cortactin signaling rewires actin-generated forces to drive basement membrane rupture and initiate breast cancer invasion

**DOI:** 10.64898/2026.04.15.717430

**Authors:** Eric Platz-Baudin, Julian Eschenbruch, Yannick Herfs, Georg Dreissen, Ronald Wein, Emiel P C van der Vorst, Michael Rose, Rudolf Merkel, Erik Noetzel

## Abstract

Oncogenic HRas activation plays a fundamental role in tumorigenesis, yet the cellular mechanisms by which HRas downstream signaling drives basement membrane (BM) disruption during early breast cancer invasion remain unclear. Using HRas-inducible breast spheroids, we demonstrate that HRas rewires cellular mechanotransduction of tumor-associated extracellular matrix stiffening to promote invasion. This process occurs independently of canonical myosin II-mediated contractility and proteolytic BM degradation. Transcriptomic and kinome profiling identified an HRas-Src-cortactin-Arp2/3 signaling axis that generates disruptive mechanical BM stress. We describe cortical triplet (CT) structures, defined by cortactin-dependent actin reinforcement and localized BM loss. CTs integrate increased cortical tension, actin polymerization forces, and myosin I-dependent contractility, thereby predicting invasion events. Pharmacological inhibition of Src or Arp2/3 reduced CT formation and invasion. Furthermore, elevated expression of HRas-cortactin-Arp2/3 axis components correlated with poor patient survival. Together, these findings uncover a previously unrecognized mechanism of early breast cancer invasion and highlight potential therapeutic targets.

## Introduction

Aberrant activation of Ras signaling drives pathological programs in various cancers ^1^. In breast cancer, Ras hyperactivation occurs through dysregulation of multiple oncogenic pathways to foster cancer progression ^2^. Cells with hyperactive Ras acquire metastatic properties ^3, 4^ and must cross the basement membrane (BM) to infiltrate the tumor microenvironment (TME). The BM scaffold acts as a cell-anchoring adhesion platform and a mechanical barrier that preserves epithelial architecture and function ^5, 6^. It consists of laminins, collagen IV, nidogen and perlecan, which together form a dense, nanoporous scaffold with substantial mechanical resistance to cell and ECM forces ^7, 8^.

Early mechanistic concepts of BM invasion focused on proteolytic degradation by matrix metalloproteinases (MMPs), typically concentrated at invadopodia ^9, 10^. However, in clinical trials MMP inhibitors neither blocked metastasis nor improved patients’ survival ^11^. More recent studies have shown that cells can apply physical forces to break through the BM barrier: actomyosin contractility acts together with proteases to weaken and disrupt the matrix ^12^. Cancer spheroids form contractile protrusions that stretch and rupture the BM ^13^. In other cases, epithelial cell clusters expand their volume and increase local contractility to compromise BM integrity ^14^. Such findings have shifted attention toward the force generating and transmitting cytoskeletal machinery as a major contributor to BM disruption. The underlying molecular mechanisms of force generation are largely based on actin-myosin interactions: Non-muscle myosin II motor protein generates high contractile forces by cooperative parallel actin filament sliding, while myosin I/V/VII mono- or dimers act on filamentous actin to produce more locally constrained actin-cell membrane tension and protrusive forces ^15, 16^. Besides myosin, cells use actin polymerization forces for actin filament and network growth. This requires various actin stabilizing factors, such as profilin or cortactin, as well as capping and geometry-shaping factors. The Arp2/3 complex is important to bundle and branch actin filaments thereby shaping the cellular cortex ^17–19^.

The mechanical properties of breast tissue undergo dramatic changes during cancer progression. Stromal collagen cross-linking and fiber realignment progressively stiffen the extracellular matrix (ECM) ^20–22^. TME stiffening steers cancer cell migration and promotes invasion through mechanosensation and -transductive signaling cascades ^23–25^. The cellular mechanoresponse to ECM stiffening involves reorganization of the actin cytoskeleton, the activation of the contractile machinery, and finally significantly increased actin forces ^26^. This mechanoadaptation enables cells to breach the BM ^12^. BM breakdown marks a critical step in the transition from ductal carcinoma *in situ* to invasive carcinoma ^27^. Despite its importance, the cellular mechanisms that control this pathological step remain incompletely defined.

To clarify these tumor-promoting processes, it is essential to understand how oncogenic signaling integrates with mechanical TME cues ^28^. Oncogene activation can alter cellular mechanosensation and force generation ^29^ to amplify oncogenic pathways ^30–32^. This bidirectional crosstalk may play a decisive role in breast cancer, where increasing matrix stiffness correlates with disease progression. However, the exact mechanisms by which HRas oncogene cooperates with altered tissue mechanics to drive BM invasion remain unclear.

Here, we established breast spheroids based on MCF10A/ER:HRAS^V^^12^ cells ^33^ to dissect the cellular events that drive early BM breach under oncogenic HRas signaling. This 3D cell model allowed temporal control of HRas hyperactivation after the formation of basoapically polarized BM-covered spheroids. We examined how HRas downstream signaling disrupts BM integrity and how this effect depends on tumor microenvironment-like matrix stiffness. We investigated a previously unrecognized oncogenic signaling axis that links HRas activity and altered ECM mechanosensation with cell invasion. We analyzed the interplay of actin cortex remodeling, myosin I- and Arp2/3-driven actin polymerization forces for mechanical BM disruption. Finally, an *in-silico* evaluation was performed to correlate prognostic implications of the HRas-cortactin-Arp2/3 -axis for breast cancer patients. Together, this study aimed to define a new mechanism by which Ras oncogene activation overrides BM barrier function and initiates early invasive transition of breast epithelial cells.

## Results

### HRas hyperactivation induces invasive transition of healthy breast cell spheroids

To monitor basement membrane (BM) disruption and cell transmigration, we generated HRas-inducible breast spheroids derived from single MCF10A/ER:HRAS V12 cells ^34, 35^ (hereafter referred as MCF10A/HRAS). These cells express an estrogen receptor (ER)-HRasV12 fusion protein that can be pharmacologically activated by the ER ligand 4-hydroxytamoxifen (OHT). After 10 days in culture (DiC) (Fig. 1A), MCF10A/HRAS spheroids (not induced with OHT) featured a similar morphology as MC10A wildtype spheroids lacking the ER:HRAS V12 modification ^5^ (Fig. 1B): In both spheroid variants, a stable apicobasal polarity was established, as indicated by inward-oriented Golgi organelles. Importantly, a continuous BM scaffold was formed and maintained that surrounded the outer basal cell layer (Fig. 1B, compare arrowheads in i and ii).

**Figure 1:**
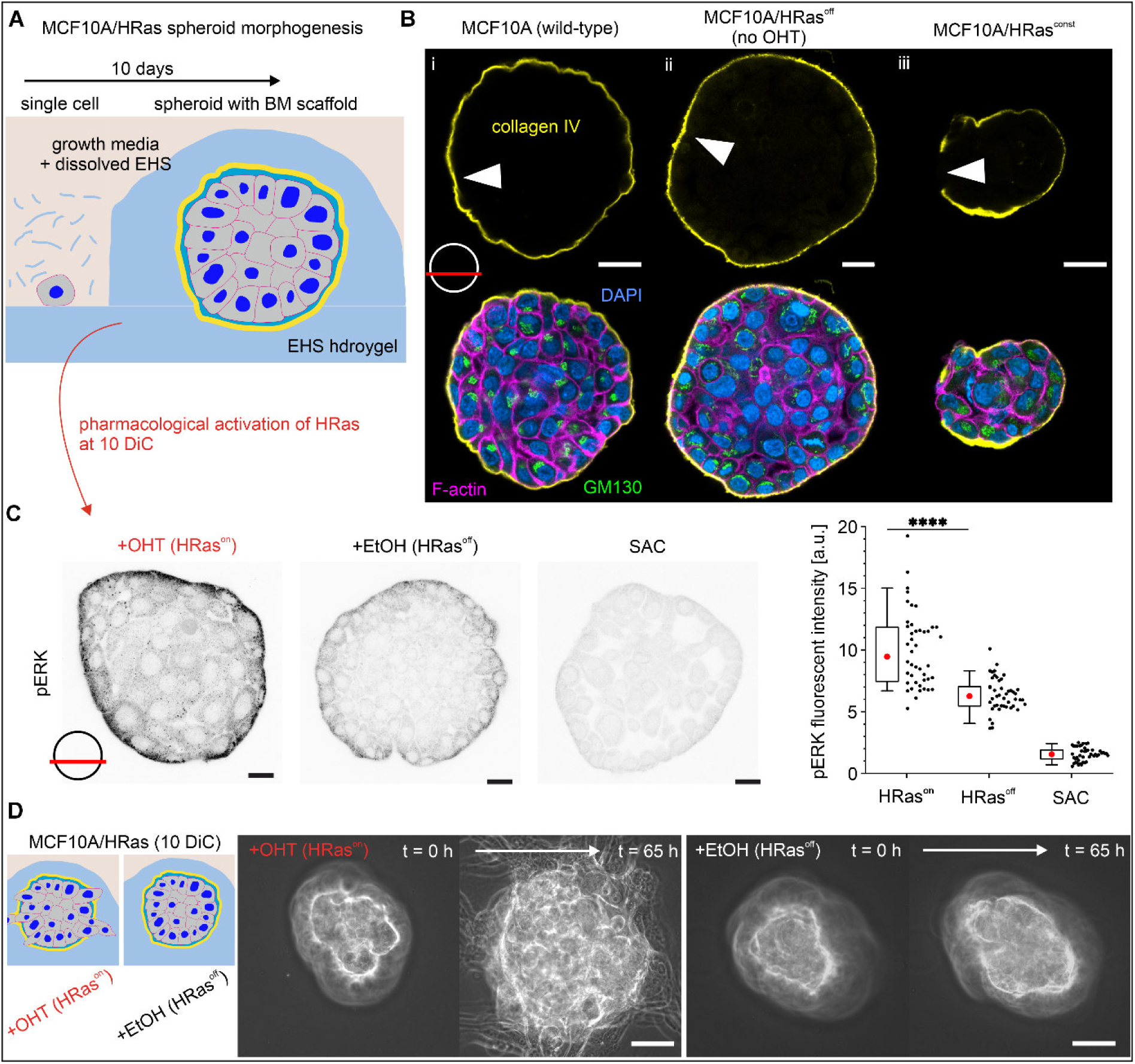
HRas hyperactivation induces invasive transition of normal breast gland spheroids. (A) Schematic of the spheroid morphogenesis assay. Single MCF10A/HRAS or MCF10A wild-type cells were cultured in a collagen IV/laminin-rich EHS(Engelbreth-Holm-Swarm) hydrogel to generate basoapically polarized spheroids after 10 days in culture (DiC). (B) Representative immunofluorescence micrographs show differences in basoapical polarization of MCF10A spheroids at 10 DiC depending on HRas activation status. BM (collagen IV, yellow), F-actin cytoskeleton (magenta), nuclei (DAPI, blue) and Golgi protein (GM130, green). (C) HRas activation confirmed by pERK immunofluorescence after 1 hour OHT or EtOH treatment. Representative immunofluorescence intensities of intracellular pERK protein (inverted grey scale) in MCF10A/HRAS spheroids treated with OHT or EtOH for 16 hours. SAC: secondary antibody control. Right, quantification of mean pERK intensity per spheroid (n ≥ 44; 3 independent experiments). Box: interquartile range; whiskers: 5th–95th percentiles; red dots: median. (D) Phase-contrast images show the invasive transition of spheroids (10 DiC) with cell transmigration into the EHS matrix after 65 hours of HRas activation with OHT. EtOH-treated HRas^off^ controls remained non-invasive. Kolmogorov-Smirnov test was performed for the data in C; n.s.: p > 0.05; ****: p ≤ 0.0001. Scale bars: 20 µm (B); 50 μm (C). Position of focal plane used for imaging and analyses is indicated by red bar.

In contrast, MCF10A/HRas^const^ spheroids, derived from cells that constantly express an active form of the HRas V12 protein, showed malfunctioned breast spheroid morphogenesis with perturbed apical cell polarization, as indicated by randomly orientated Golgi protein. More importantly, HRas^const^ led to discontinuous BM scaffolds (Fig. 1B, arrowhead in iii). This clearly demonstrated that HRas hyperactivation during early stages of breast gland development disrupts basoapical polarization and BM formation. Consistent results of misshaped HRas-activated MCF10A spheroids have been reported ^36^. Thus, it implicated the necessity to induce HRas in spheroids that gained proper polarity and BM scaffolds to analyze its oncogenic effects.

For this purpose, we made use of temporally controlled HRas activation in mature MCF10A/HRas spheroids featuring fully developed BMs (10 DiC). At this point OHT treatment (16 hours) reproducibly induced phosphorylation of the HRas downstream effector ERK ^35^ (Fig. 1C). Quantification of pERK intensity levels revealed a significant increase of ≈60% in OHT-treated spheroids compared to the EtOH control (Fig. 1C). The pERK increase was most pronounced in the basal cell layer facing the BM-ECM interface. This activation gradient was most likely caused by decreased drug accessibility towards the spheroid center. In the following, OHT-treated spheroids are referred to as HRas^on^ spheroids and the untreated EtOH controls as HRas^off^ spheroids. Further, HRas activation frequently led to the invasive cell transition of non-transformed breast spheroids that normally remain in a homeostatic state: After 65 hours of HRas activation, 43% of spheroids showed invasive cell streams that breached the BM reaching into the microenvironment. In contrast, HRas^off^ controls remained polarized without any visible morphological changes (Fig. 1D).

These results show that HRas hyperactivation alone induced an invasive transition in originally non-transformed breast epithelial spheroids, even under microenvironmental conditions that reflect the compliance of healthy breast tissue ^37, 38^. This spheroid model thus provided a controlled and reproducible platform to investigate the functional consequences of HRas activation *in vitro*.

### HRas invasion bypasses tumor stiffness sensing, myosin II forces and BM proteolysis

After showing the pro-invasive consequences of HRas hyperactivation, we examined whether tumor-like ECM stiffness, as fundamental mechanical cue, could enhance HRas invasion. Previous work has shown that increased ECM rigidity promotes BM rupture in non-transformed breast epithelial spheroids ^5^. We therefore established a quantitative assay (Fig. 2A) to measure HRas-induced BM transmigration and cell invasion (Fig. 2B - C) as a function of microenvironmental stiffening (see Supplementary Video 1).

**Figure 2:**
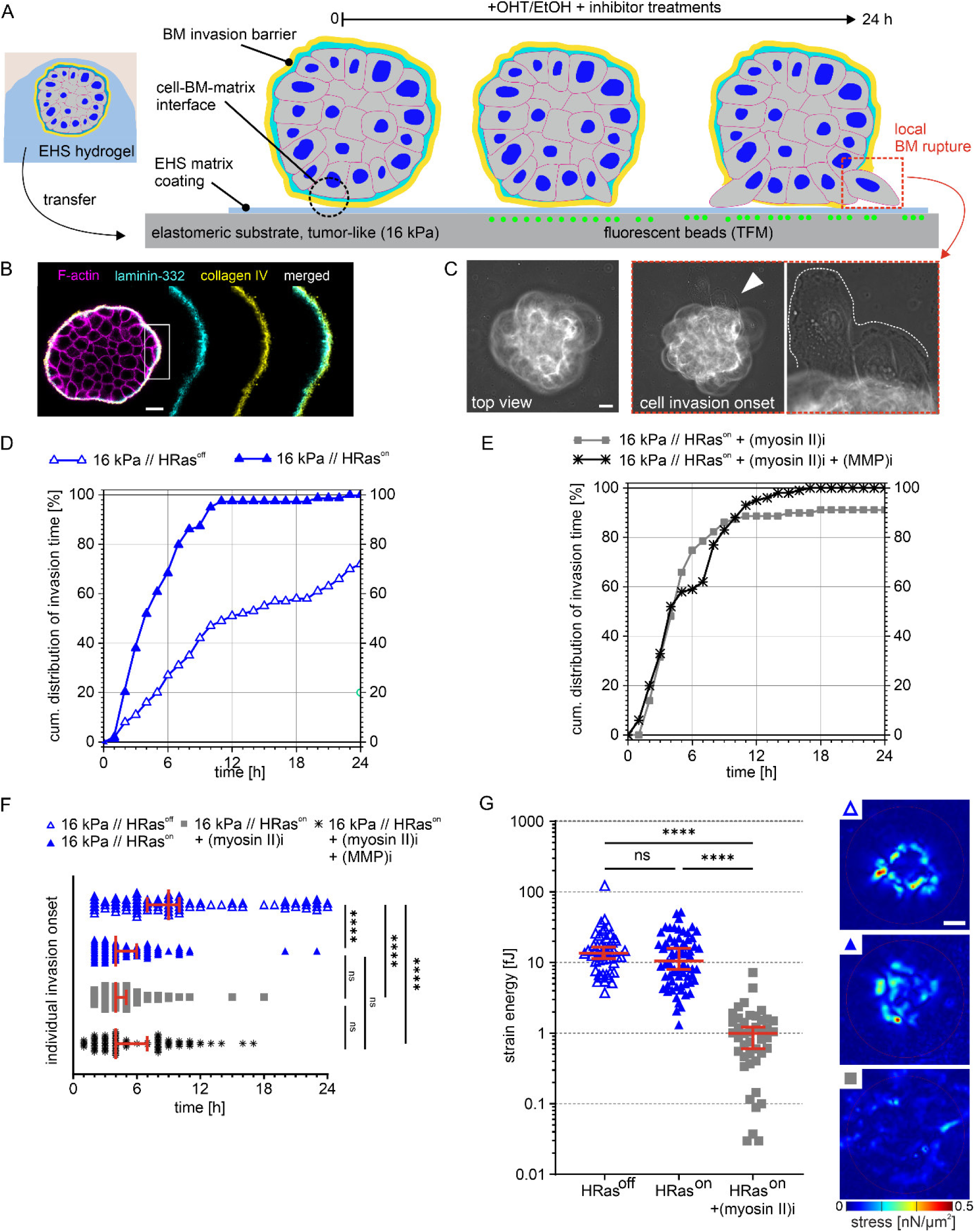
HRas-driven invasion bypasses tumor stiffness sensing, myosin II forces and BM-proteolysis. (A) Scheme of BM disruption and cell invasion assay. MCF10A/HRAS spheroids (10 DiC) were isolated from EHS matrix and placed on elastomeric substrates (16 kPa, functionalized with EHS proteins) to count events of local BM rupture and cell transmigration. Mechanical BM stress exertion by breast spheroids at time point of invasion onset was measured by traction force microscopy (TFM): Surface-coupled fluorescent fiducial microbeads were used to track tangential surface deformations from which strain energies were calculated as measure for cell force-generated BM stress. (B) In spheroids, the outer basal cell layer is covered by a BM which itself is in contact to the underlying substrate. Images show the BM integrity of a representative HRas^on^ sample, fixed and stained after adhering (1 hour) to the elastomeric substrate. Collagen IV (yellow), laminin-332 (cyan) and F-actin cytoskeleton (magenta). Zoom in highlights *in vivo*-like layering of the endogenous BM. (C) Representative sequence of phase-contrast images illustrates the first appearance of protrusive cell bodies (also shown as zoom in), marking onset of BM disruption and cell transmigration. This was counted as a positive event of invasion. (D) Cumulative distribution of BM disruption time, depending on HRas induction on 16 kPa substrates (n ≥ 79 spheroids of ≥ 3 independent experiments). (E) Cumulative distribution of BM disruption time in spheroids treated with blebbistatin for myosin II inhibition and additionally with marimastat for MMP inhibition after HRas induction on stiff 16 kPa substrates (n ≥ 69 spheroids of ≥ 3 independent experiments). (F) Scatter plot shows individual invasion onset time points for the sample conditions analyzed in (D and E) (median and 95% confidence interval (CI)). (G) Calculated strain energies (SE) exerted by individual spheroids at onsets of BM disruption, depending on HRas activation and actomyosin inhibition, (cf. D and E). Representative maps of cell-induced traction stresses per condition from which SE were calculated. Scatter plot: median with 95% CI (n ≥ 48 from 3 independent experiments). Kruskal-Wallis test with Dunn’s multiple comparison test was performed for the data in D and E; n.s.: p > 0.05; *: p ≤ 0.05; **: p ≤ 0.01; ***: p ≤ 0.001; ****: p ≤ 0.0001. Scale bars: 20 µm (B, C and G).

HRas^off^ spheroids cultured on tumor-like stiff substrates exhibited pronounced invasion (72% within 24 hours; median onset: 9 hours; Fig. 2D and F). In contrast, same spheroid group on physiological normal-like ECM stiffness did not undergo invasion (cf. Fig. 1D). These results demonstrate that HRas^off^ spheroids are highly mechanosensitive and undergo invasion in response to TME stiffening. Notably, the standard growth medium used in these initial experiments induced basal activation of the ER:HRas fusion protein, thereby enhancing invasion. This residual activation was attributable to estrogenic phenolic compounds (see Supplementary Fig. S1). Consistently, MCF10A spheroids lacking the ER:HRas construct displayed lower invasion under comparable conditions ^5^. To eliminate this interfering effect, subsequent experiments were performed in steroid hormone-free conditions, which reduced the TME stiffness-dependent invasion to 50% (median onset: 4.2 hours). This had no significant impact on HRas^on^ spheroids (median onset: 9.5 hours). These data demonstrated that oncogenic HRas is sufficient to drive invasive cell transition independently of TME stiffening, while strongly synergizing with extracellular mechanical cues.

We asked whether HRas^on^ spheroids convert TME stiffness into altered cellular forces that could explain the rapid invasive cell transition. Because actomyosin contractility is a key driver of BM stress and rupture ^23^, we inhibited non-muscle myosin II-ATPase activity. This treatment did not affect the invasion outcome (cum. frequency: 100%, median onset: 5 hours; Fig. 2C and D). To quantify cell-derived BM stress, we performed traction force microscopy (TFM) and calculated strain energy (SE) as a readout of cellular traction transmitted through the BM onto the underlying substrate ^5^. At the time of BM rupture, HRas^off^ and HRas^on^ spheroids generated comparable SE (13.6 fJ, IQR: 8.2–20.2 fJ vs. 10.5 fJ, 5.5–19.3 fJ; Fig. 2E). Myosin II inhibition reduced SE to near-background levels (1.0 fJ, 0.5–1.5 fJ), which confirmed effective suppression of contractility by blebbistatin drug. Under these conditions, SE values were close to background levels measured on cell-free substrates (Supplementary Fig. S2). These findings demonstrated that HRas-driven invasion occurs without elevated actomyosin-mediated cell contractions that could account for disruptive BM-stress.

Finally, we tested whether proteolytic BM degradation contributes to invasion by co-inhibition of myosin II and major invasion-relevant MMPs (MT1-MMP, MMP-1, -2, -3, -7, -9) using the broad-spectrum inhibitor marimastat ^39^. MMP inhibition did not affect invasion (cf. Fig. 2C and D). Consistently, immunostaining showed comparable MT1-MMP localization and signal intensity in the basal cell layer of both HRas^on/off^ spheroids (Supplementary Fig. S3). Together, these data indicate that HRas-driven invasion is independent of MMP-mediated BM degradation.

### HRas triggers BM disruption through local actin cortex thickening and reinforcement

Neither inhibition of myosin II nor combined blockade of MMPs reduced the invasiveness of HRas^on^ spheroids, despite the clear lack of detectable pro-invasive contractile forces. This indicated alternative, non-canonical force-generating mechanisms. We therefore examined the F-actin cytoskeleton organization within the basal cell layer at the cell-BM-substrate (CBS) interface where BM rupture occurred: Immunostaining revealed sequential cytoskeletal changes after HRas induction (Fig. 3A). Within the first 1.5 hours, pre-invasive spheroids had high coverage of actin-rich microspikes (MS) that were oriented towards the substrate and previously described as ECM stiffness-sensing units ^23^. After 4.5 hours, cortical actin networks thickened at BM contact sites. Figure 3A highlights such cortex thickening accompanied by laterally oriented MS bundles (see i, arrow heads). After 5 hours, these regions showed cortical actin reinforcement that frequently coincided with local BM scaffold loss (Fig. 3A, ii arrowheads). Figure 3B shows a detailed view on late-stage invasion characterized by local BM rupture (i) and cell transmigration into the microenvironment (ii + iii).

**Figure 3:**
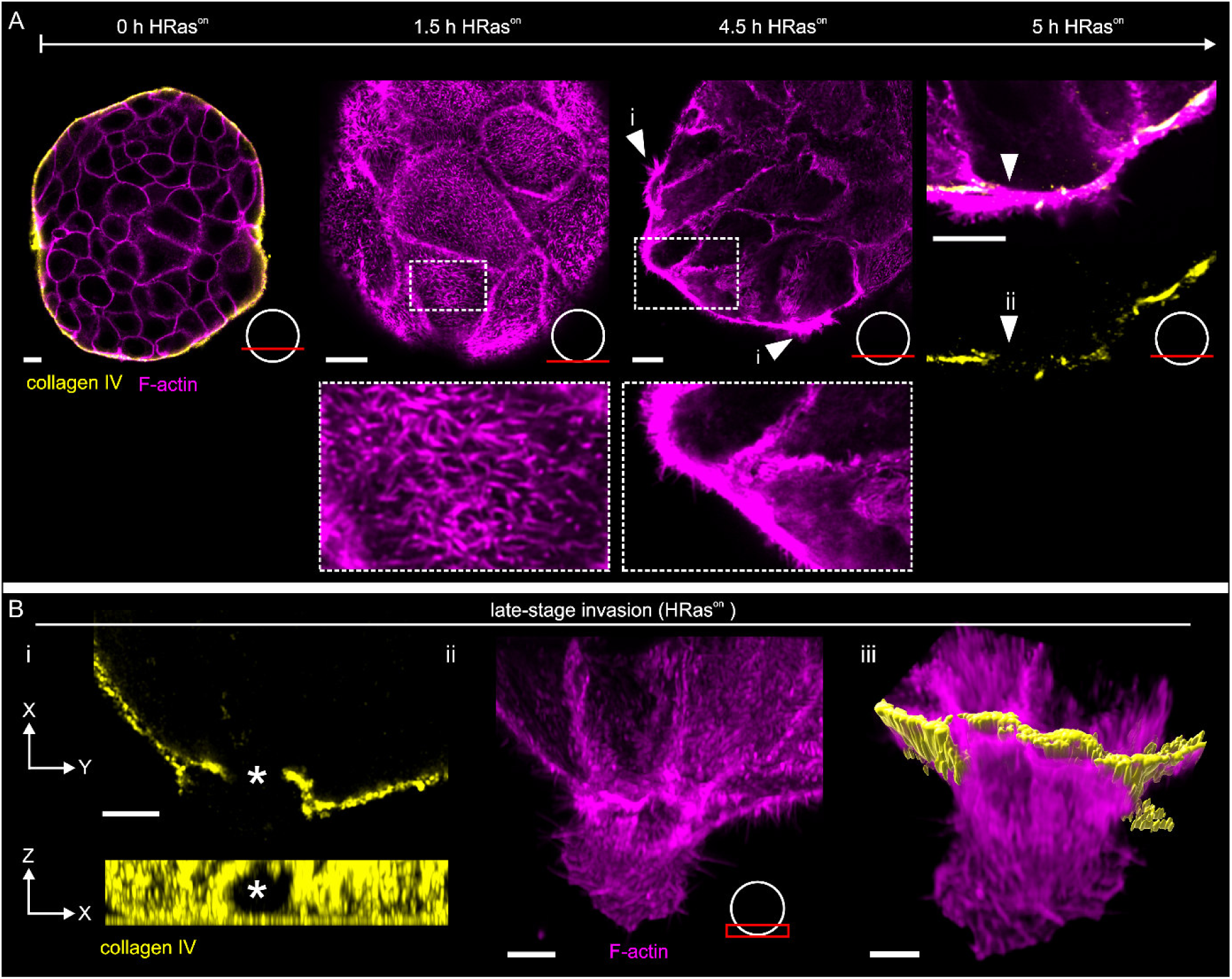
HRas hyperactivation triggers BM disruption through local actin cortex reinforcement. (A) Time-resolved immunofluorescence staining reveals cytoskeletal remodeling following HRas activation. Spheroids were stained for F-actin (magenta) and BM (collagen IV, yellow) and imaged at the CBS-interface using 16 kPa elastomeric cell adhesion substrates. Pre-invasive HRas^on^ spheroids (1.5 h and 4.5 h) showed dense MS (i, arrowheads). After 5 hours, pronounced cortical actin reinforcement appeared at sites of local BM loss (ii, arrowheads). (B) Representative images highlight a late-stage event of BM cell transmigration: BM-collagen IV XZ projection of an image stack shows a cell transmigration hole within the BM scaffold (i, asterisks). MIP (ii) and a 3D image stack reconstruction show the F-actin cytoskeleton of a BM-transmigrating cell (iii). Scale bars: 10 µm (A) and 5 µm (B). Position of focal plane(s) used for imaging and analyses is indicated by red bar/rectangle.

Of note, invasive cells lacked prominent F-actin stress fiber (SF) formation, which typically characterize highly contractile and BM-stressing cells ^23^. This lack of SFs was consistent with the low SE values measured before (cf. Fig. 2G). Ultimately, at the late stage of transmigration into the ECM, spheroid cells showed pronounced actin thickening at BM rupture sites that were accompanied by BM widening and hole formation (Fig. 3B, panels i and iii). These findings identified a HRas-driven program of cortical actin reinforcement that coincided with, and likely contributed to, non-proteolytic BM rupture and invasive cell transition.

### HRas rewires actin cortex dynamics to mechanically stress the BM scaffold for invasion

After finding the spatial coincidence of cortical actin thickening and BM rupture, we next analyzed how this characteristic cytoskeletal remodeling could be functionally linked with invasion progression. Using 4D LCI, F-actin and BM dynamics were resolved in high spatiotemporal manner after HRas activation.

Within the first four hours, HRas^on^ spheroids exhibited a dynamic reshaping: basal cells at the CBS-interface showed repeated pushing and pulling displacements of cortical actin patches mechanically coupled to the BM. As representative example, a single cell retracted approx. 10 µm toward the spheroid center, while reinforcing the actin cortex (Fig. 4A, arrowheads). This retraction phase was followed by local BM disruption (t = 5.5 hours). These qualitative observations link the displacement of reinforced cortex regions to local mechanical deformation and subsequent rupture of the BM cell migration barrier (see also Supplementary Video S2).

**Figure 4:**
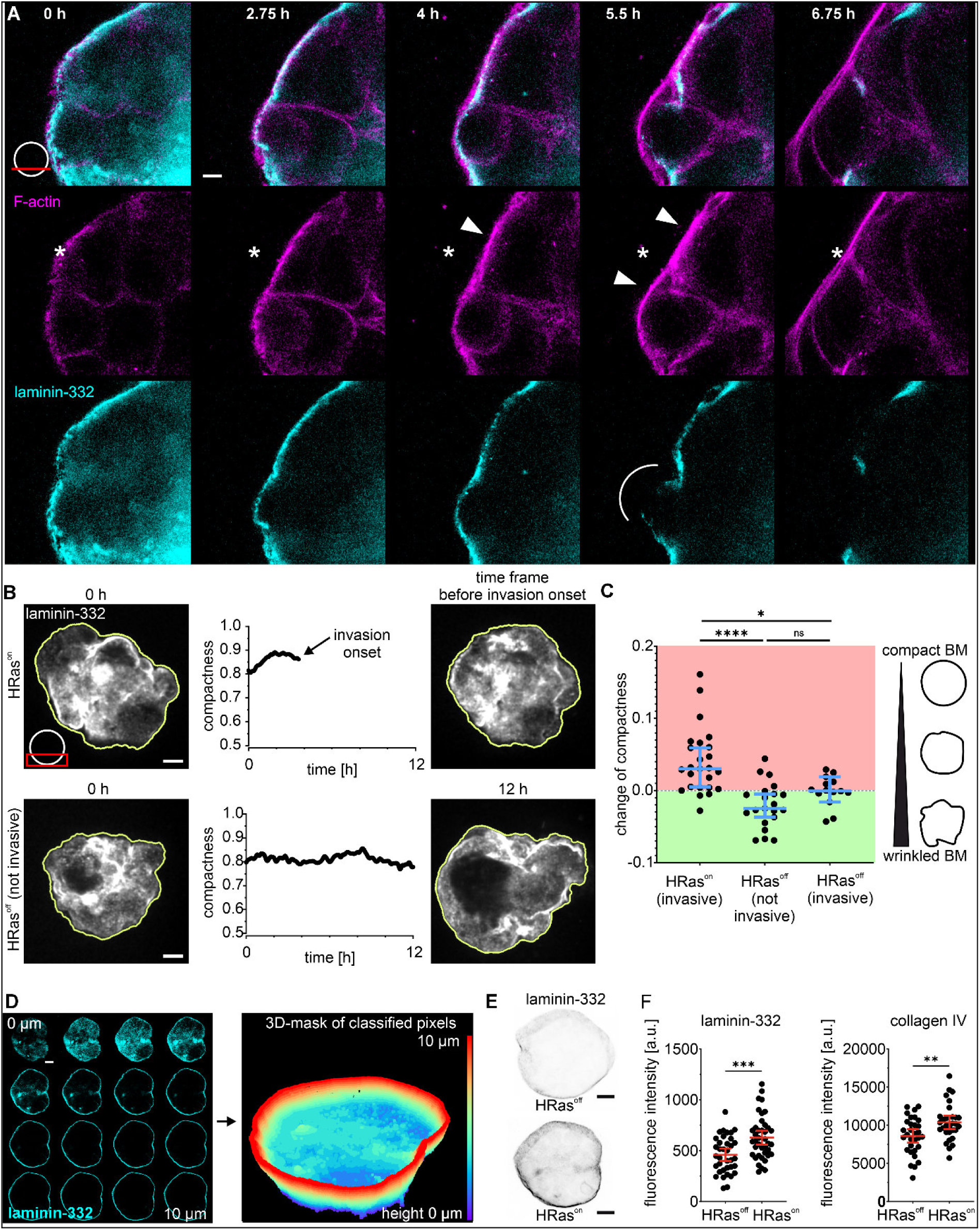
Ras activates cellular cortical actin dynamics to mechanically stress the BM scaffold. (A) Time-lapse series of HRas^on^ spheroid motion on a 16 kPa substrate. The white asterisk marks a retractive actin cortex movement; white arrowheads indicate reinforced cortical actin patches. Local BM disruption occurred at 5.5 hours after HRas activation (white line). BM is visualized with Alexa Fluor 488-conjugated laminin-332 antibody; F-actin is labeled with SiR-actin live-cell dye. Of note, a slight channel shift was caused by cell movements during image acquisition without affecting data interpretation. For complete image series see Supplementary Video 2. (B) Time-course quantification (0 – 12 hours) of spheroid compactness at the cell–BM–matrix interface. MIPs of confocal image stacks (laminin-332 signal) were used for shape analysis in HRas^on^ and HRas^off^ groups. (C) Scatter plot compares the changes of compactness for HRas^on^ and HRas^off^ spheroids (t = 0 hours vs. t = individual time points of invasion onset) and for the non-invasive fraction of HRas^on^ samples (t = 0 hour vs. t= 12 hours). (n ≥ 15 spheroids from ≥ 3 experiments). Bars and whiskers: median and 95% CI. The scheme illustrates the observed increase (smooth BM) or decrease (wrinkled BM) of spheroids compactness. (D) Representative confocal stack used to reconstruct a 3D BM shell (10 µm height) from laminin-332 signals. Within the volume of this 3D mask, signal intensities of laminin-332 and collagen IV were quantified (see Materials and Methods for detailed information). (E) Top-view on 3D-masked BM intensities (of laminin-332) in representative HRas^on^ versus control spheroids. (F) Scatter plots summarize the quantitative signal intensities BM protein intensities (laminin-332 and collagen IV) after 2 hours OHT-treatment (n ≥ 30 spheroids from ≥2 independent experiments; bars and whiskers: mean and 95% CI). Kruskal-Wallis test with Dunn’s multiple comparison was performed for data in C. Shapiro–Wilk test of normality and unpaired t-test was performed for data in F; n.s.: p > 0.05; *: p ≤ 0.05; **: p ≤ 0.01; ***: p ≤ 0.001; ****: p ≤ 0.0001. Scale bars: 5 µm (A); 20 µm (B, D, E). Position of focal plane(s) used for imaging and analyses is indicated by red bar/rectangle.

To quantify this HRas-induced BM deformation, an automated image-based BM segmentation tool was developed (Fig. 4B). The analysis used laminin-332 fluorescence to define spheroid outlines (masks) and calculate their compactness: spheroid compactness was defined as the ratio of actual area to the area of a circle with identical perimeter. In other words, compactness increases when spheroids adopt a more circular shape with smooth boundaries and decreases with irregular, wrinkled deformations (see cartoon, Fig. 4C).

HRas activation caused a progressive and significant increase in compactness, reaching a maximum prior to invasion onset (mean t = 4 hours). In contrast, non-invasive HRas^off^ spheroids maintained constant compactness over the entire time of analyses (t = 12 hours) (Fig. 4B). Further stratification of HRas^off^ spheroids into non-invasive and invasive groups revealed that exclusively HRas^on^ spheroids exhibited increased compactness (Fig. 4C) whereas the HRas^off^ group exhibited only marginal changes (t = 4 hours, range: -0.05 to +0.05) Supplementary Fig. S4A). This trend held true when measuring points of HRas^on^ and non-invasive HRas^off^ spheroid were normalized to relative time points (1/4, 2/4, 3/4 and 1): The compactness elevated only in HRas^on^ spheroids prior to invasion but not in HRas^off^ controls (Supplementary Fig. S4B).

We next tested whether the BM barrier itself was altered by the tension resulting from increased compactness in HRas^on^ spheroids. The mean fluorescence intensity of laminin-332 and collagen IV were measured as proxies for BM overall protein density. Laminin-332 signals were used to construct a 3D mask for the BM shell at the CBS-interface (Fig. 4D). Quantification within this voxel mask showed significantly elevated intensities for both BM components in HRas^on^ spheroids after 2 hours of activation. Laminin-332 increased by 37% (HRas^off^: 460, s.d. 180 a.u.; HRas^on^: 630, s.d. 220 a.u.), and collagen IV by 21% (HRas^off^: 8,600, s.d. 2.500 a.u.; HRas^on^: 10.400, s.d. 2.400 a.u.) (Fig. 4E and F). These data indicated a densification of the BM scaffold.

These findings demonstrated that increased compactness and BM densification are specific morphological adaptation of invasive HRas^on^ spheroids. They imply a functional link between HRas activation, cortical actin reinforcement, and coordinated cell movements that significantly compact both the spheroid cytoskeleton and the BM scaffold prior to cell transmigration. This process most likely generated mechanical BM stress that contributes to its disruption and facilitating invasive transition.

### HRas–Src–cortactin signaling drives cytoskeletal remodeling and BM invasion

To identify signaling events linking oncogenic HRas activity to cytoskeletal reorganization and BM disruption, we profiled transcript levels and kinase activity downstream of HRas. A targeted RT-qPCR array was used to assess cytoskeleton-related gene expression. Among the analyzed transcripts, CDK5R1 (p35) and cortactin (CTTN) were upregulated (1.35-and 1.42-fold, respectively, Fig. 5A; Supplementary Table 1). Despite higher statistical significance for CDK5R1, we prioritized cortactin based on its established role in actin organization ^40^, and its consistency with the observed HRas-induced cytoskeletal phenotype. Because cortactin activity is regulated by phosphorylation ^41^, we next assessed upstream kinase activation to identify potential regulators of cortactin in HRas^on^ spheroids.

**Figure 5:**
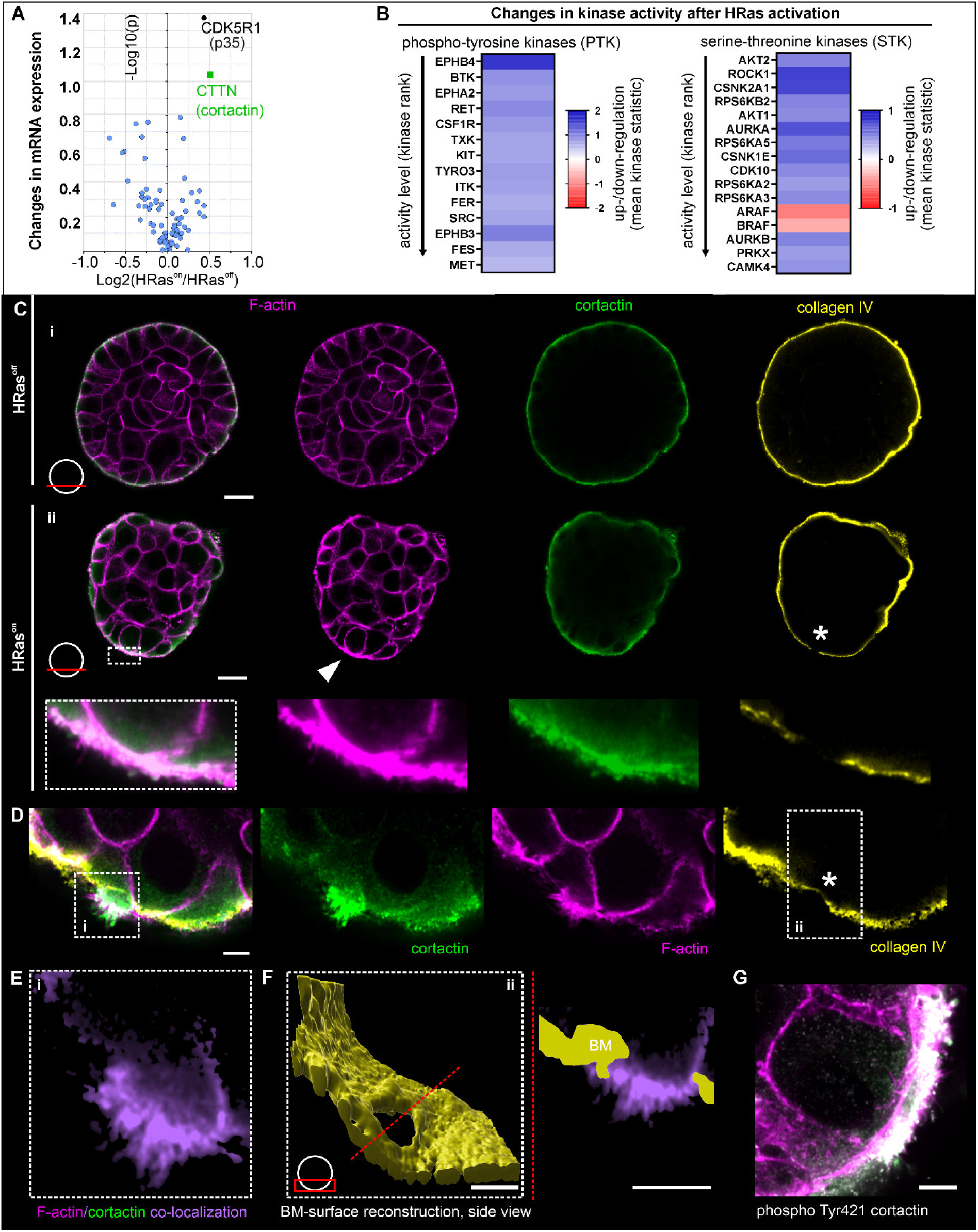
HRas rewires Src and cortactin signaling to reinforce the actin cortex at BM rupture sites. (A) Volcano plot compared the expression of 84 genes related to cytoskeletal regulation measured with a qRT-PCR array. Gene expression was analyzed after 4.5 hours of HRas activation on tumor stiffness (16 kPa). (B) Profiling analysis of phospho-tyrosine-kinase (RTK) and serine-threonine-kinase (STK) activation depending on HRas activation retrieved from kinase-substrate microarrays. Kinase rank: decreasing p-values of change in kinase activity. Mean kinase statistic: color coded up (1)- and down (-1)-regulation. Kinase regulation was compared after 1 hour of OHT to EtOH treatment (16 kPa substrates). (C) Immunostained and fixed spheroids to compare the actin cytoskeleton (F-actin, magenta), cortactin (green) and BM scaffold (collagen IV, yellow,) depending on HRas activation. White arrowhead indicates reinforced cortical actin with cortactin accumulation; white asterisk marks a site of BM rupture. (D) Detailed view of a HRas^on^ spheroid highlights a patch of reinforced actin, cortactin incorporation and MS protrusion bundles within a BM tunnel for cell transmigration. The white asterisk marks the ruptured BM site. (E) Image shows blue-outlined inset in (D) of actin-cortactin-protrusion (purple). Co-localization was calculated in Imaris (see Materials and Methods for detailed information). (F) BM surface reconstruction (yellow) covering the orange-outlined inset in (D). On the right, top view along dashed line on the protrusion bundle through the BM hole. (G) Representative image of pronounced pTyr421 localization within the thickened actin cortex areas. (See Supplementary Fig. S6 for full image series with single channel display). Scale bars: 20 µm (C), 5 µm (D, F and G). Position of focal plane(s) used for imaging and analyses is indicated by red bar/rectangle.

We profiled phosphotyrosine (PTK) and serine/threonine kinase (STK) activities using a peptide-based microarray. Several kinases were differentially regulated in HRas^on^ spheroids compared to controls (Fig. 5B; Supplementary Table 2). EPHB4 and AURKA emerged as candidate mediators of HRas-dependent invasion ^42–45^. However, inhibition of either kinase did not affect invasion, with HRas^on^ spheroids maintained their rapid invasion outcome (median onset: 5 hours for EPHB4 and 4 hours for AURKA inhibition, Supplementary Fig.S5).

Integrated transcriptome and kinome analyses revealed increased Src expression and activity. Src is a known effector of oncogenic Ras signaling ^46–48^ and regulates cortactin phosphorylation ^41^. We therefore examined the HRas–Src–cortactin axis in detail: Cortactin protein levels and localization were compared between HRas^on^ and control spheroids. In both conditions, cortactin localized predominantly to the basal cell layer within cortical actin networks adjacent to the BM (Fig. 5C i and ii). In HRas^on^ spheroids, cortactin accumulated in reinforced cortical actin patches (Fig. 5C ii, white arrowhead).

Strikingly, these patches localized adjacent to sites of BM disruption (Fig. 5C ii, asterisk). In contrast, HRas^off^ spheroids lacked this co-localization pattern and BM damage (Fig. 5C i). To assess Src-dependent cortactin activation, we stained for pTyr421-cortactin ^41^: pTyr421-cortactin co-localized with thickened cortical actin (Fig. 6G). These data link Src activation to cortactin phosphorylation at reinforced cortical actin sites. At later stages, HRas^on^ cells formed prominent bundles of actin MS (cf. Fig. 2F). These protrusions penetrated the BM, forming transmigration paths (Fig. 5D, asterisk). Cortactin was enriched within these BM-penetrating structures (Fig. 5E and F).

**Figure 6:**
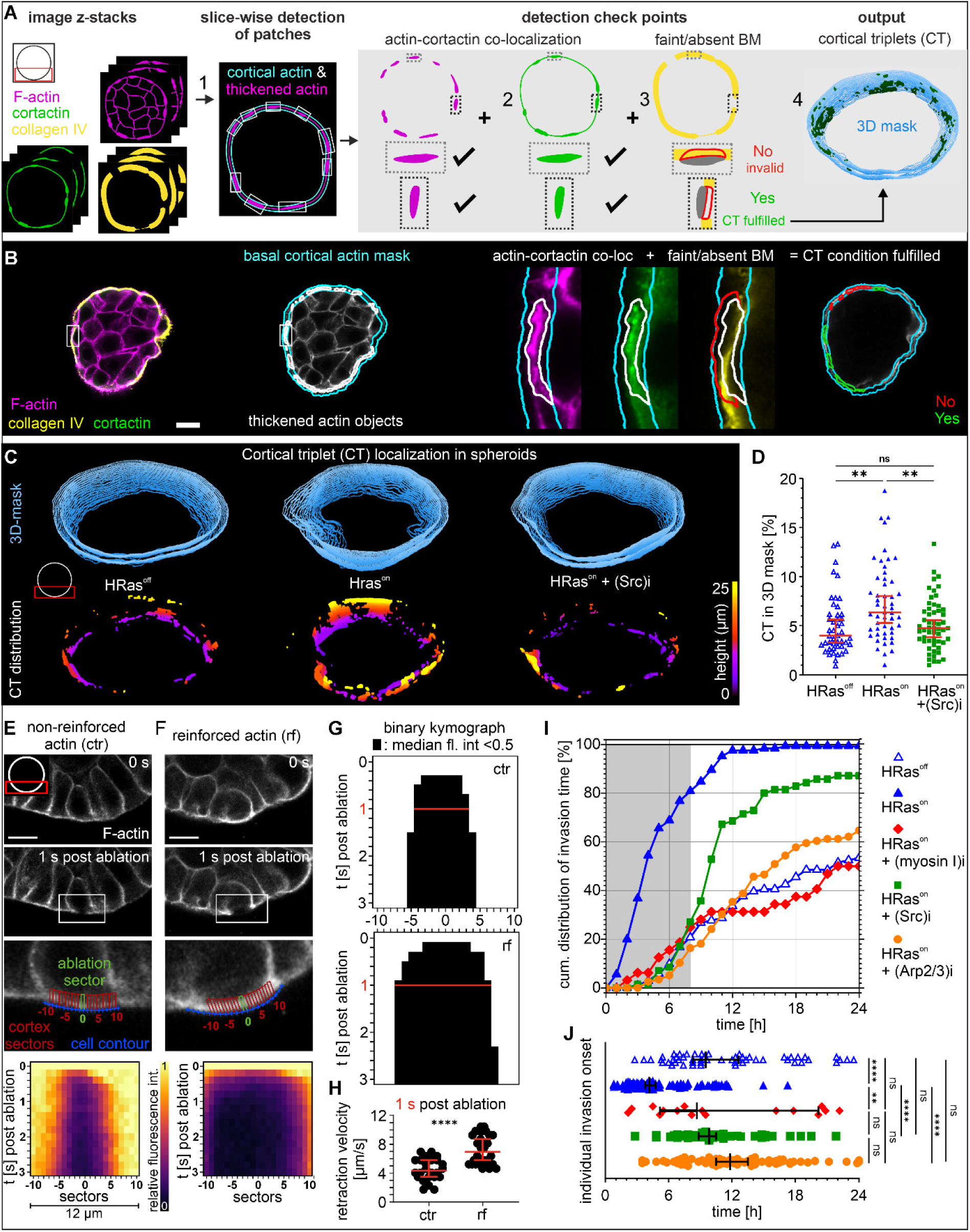
Cortical triplets and myosin I are functionally linked with BM rupture and cell invasion. (A) Schematic workflow illustrates the automatic detection of cortical triplets (CTs) within the basal cell layer of spheroids (fixed and stained after 3.5 hours of OHT or EtOH treatment, 16 kPa substrate): Sequential analyses of Confocal image z-stacks (0) for the (1) detection of reinforced cortical actin sites being (2) co-localized with cortactin signal. Co-localization negative objects were excluded. (3) Positive evaluated objects were checked for co-appearance of faint/absent BM signal. Red outline: co-localization mask. (4) Objects that fulfilled all three criteria were classified as CT. 3D masks serve to calculate the overall CT coverage in a BM volume of 25 µm height. (B) Micrographs of a representative spheroid show steps 1 to 4 of the CT detection routine (for detailed information on object detection see Supplemental Methods). (C) Representative 3D masks show CT-coverage of the basal cell layer at the CBS-interface, depending on HRas activation and Src inhibition. (D) Scatter plot summarizes the CT-coverage of individual spheroids (n ≥ 46 for each sample condition, from 3 independent experiments) displayed with median and 95% CI. Representative cortex laser ablation of HRas^on^ spheroids at non-reinforced control sites (E) and CT-associated reinforced actin sites (F). Actin loss was monitored in 12 µm sectors centered on the ablation site. Kymographs show fluorescence intensity normalized to pre-ablation levels. A decrease in relative actin fluorescence intensity <0.5 was defined as rupture. (G) Binary kymographs display sectors with median relative fluorescence intensity <0.5 (black; n = 24 control and n = 30 reinforced sites from 3 independent experiments). Red lines indicate median rupture width at 1 s post-ablation; corresponding retraction velocities are shown in (H) (scatter with median ± 95% CI). (I) Cumulative distribution of BM disruption time, depending on HRas activation and combined with myosin I, Src kinase or Arp2/3 inhibition (n ≥ 33 spheroids, per condition from ≥ 2 independent experiments). (J) Individual events of BM disruption onset over time for the samples analyzed in (I) displayed with median and 95% CI. Kruskal-Wallis test with Dunn’s multiple comparison test was performed for the data in (D and F); Mann-Whitney test was performed for the data in (H): n.s.: p > 0.05; *: p ≤ 0.05; **: p ≤ 0.01; ***: p ≤ 0.001; ****: p ≤ 0.0001. Scale bars: 20 µm. Position of focal planes used for imaging and analyses is indicated by red rectangle.

Together, these findings revealed a spatiotemporal coupling of HRas activation, pTyr421-cortactin–mediated actin reinforcement, and BM disruption.

### HRas drives cortical triplet formation and myosin I-mediated BM rupture and invasion

HRas induced co-localization of reinforced cortical actin, phosphorylated cortactin, and local BM rupture. We termed this pattern “cortical triplet” (CT). We developed an image analysis pipeline with detection checkpoints to quantify CT formation and classify HRas-driven invasion events (Fig. 6A and B). This analysis yielded CT coverage across the basal cell layer (see 3D masks, Fig. 6A 4).

CT coverage was significantly increased in HRas^on^ spheroids compared to controls (Fig. 6C). Given that Src phosphorylates cortactin at Tyr421 to promote actin assembly ^49^ (see also Fig. 5G), we inhibited Src activity. Src inhibition (PP2) reduced CT formation to control levels (Fig. 6C). Quantitative analysis revealed a 1.6-fold increase in CT coverage in HRas^on^ spheroids, which was abolished upon Src inhibition (Fig. 6D). Consistently, Src inhibition reduced both actin reinforcement and cortactin co-localization events (Supplementary Fig. S7).

We next tested whether reinforced cortical actin patches alter cortical tension to drive BM rupture. To this end, reinforced cortical actin regions in living HRas^on^ spheroids were locally cut using laser-assisted nanosurgery. Local cortical tension was quantified by comparing ablation-induced tangential retraction of actin fluorescence between thickened and thin cortical regions (Fig. 6E and F). Reinforced regions retracted faster and over greater distances than control regions (Fig. 6E; see also Supplemental Videos S3 and S4). Accordingly, reinforced regions exhibited more pronounced ruptures (Fig. 6G). At 1 s post-ablation, retraction velocity was 1.9-fold higher in reinforced regions, with 50% of values exceeding the maximum observed in control (Fig. 6H and 6G, red lines). Faster actin retraction has been previously shown to indicate elevated local cortical tension ^50^, which was also evident at reinforced actin sites in HRas-induced spheroids.

Consistent with elevated cortical tension, we inhibited myosin I activity in HRas^on^ spheroids. Myosin I links cortical actin to the plasma membrane ^51^ and may transmit disruptive forces to the BM. Myosin I inhibition markedly reduced invasion (50%; median onset: 9 hours), approaching HRas^off^ levels (Fig. 6I and J).

We next tested whether CT formation directly contributes to invasion. Src inhibition (8 hours) reduced invasion by >60% and delayed onset (∼11 hours), comparable to HRas^off^ controls (Fig. 6I, grey box and J). However, overall invasion incidence remained partially reduced after 24 hours (87%).

Cortactin co-localizes with CTs (Fig. 5D) and promotes Arp2/3-mediated actin polymerization ^52, 53^. To assess the role of actin polymerization, we inhibited the Arp2/3 complex. Arp2/3 inhibition attenuated invasion, with delayed onset to 12 hours and reduced incidence to 65% after 24 hours, similar to HRas^off^ controls.

Together, these data identify Src- and cortactin-dependent Arp2/3-mediated actin polymerization, coupled to myosin I, as the primary mechanical driver of BM disruption in HRas-activated spheroids, explaining the negligible impact of myosin II inhibition

### HRas–cortactin–Arp2/3 axis predicts poor outcome in HRas-activated breast cancer

Given the strong link between cortactin overexpression and phosphorylation, its cellular redistribution, and HRas-driven invasion, we evaluated the clinical relevance of this signaling axis using breast cancer data from “The Cancer Genome Atlas (TCGA, National Institute of Health)”. We stratified tumors by HRAS mutations and copy number variation, either “wild-type / loss” (HRas^-/-^, n=667) or “mutated / amplified” (HRas^+/+^, n=95). The latter was indicative of elevated HRas activity in HRas^on^ spheroids. In the HRAS^+/+^ subgroup, high cortactin (CTTN) gene expression correlated with worse recurrence-free survival (RFS) (Fig. 7B). In contrast, in the HRas^-/-^ subgroup, elevated CTTN levels were associated with favorable RFS (Fig. 7A). Applying the same stratification to ARP2/3 complex subunits revealed that high expression of the three subunits ARPC1B, ARPC2 and ARPC4 significantly predicted poorer RFS in HRas^+/+^ tumors, whereas no adverse effect appeared in the HRas^-/-^ group (Supplementary Fig. S8). Since the functional Arp2/3 complex requires all of its seven subunits ^54, 55^, these findings point to Arp2/3-mediated actin polymerization within the cell cortex as a key driver of invasiveness in HRas hyperactivated breast cancer. Collectively, the data highlighted a clinically relevant role for the HRas–cortactin–Arp2/3 axis in breast cancer invasion and prognosis.

**Figure 7:**
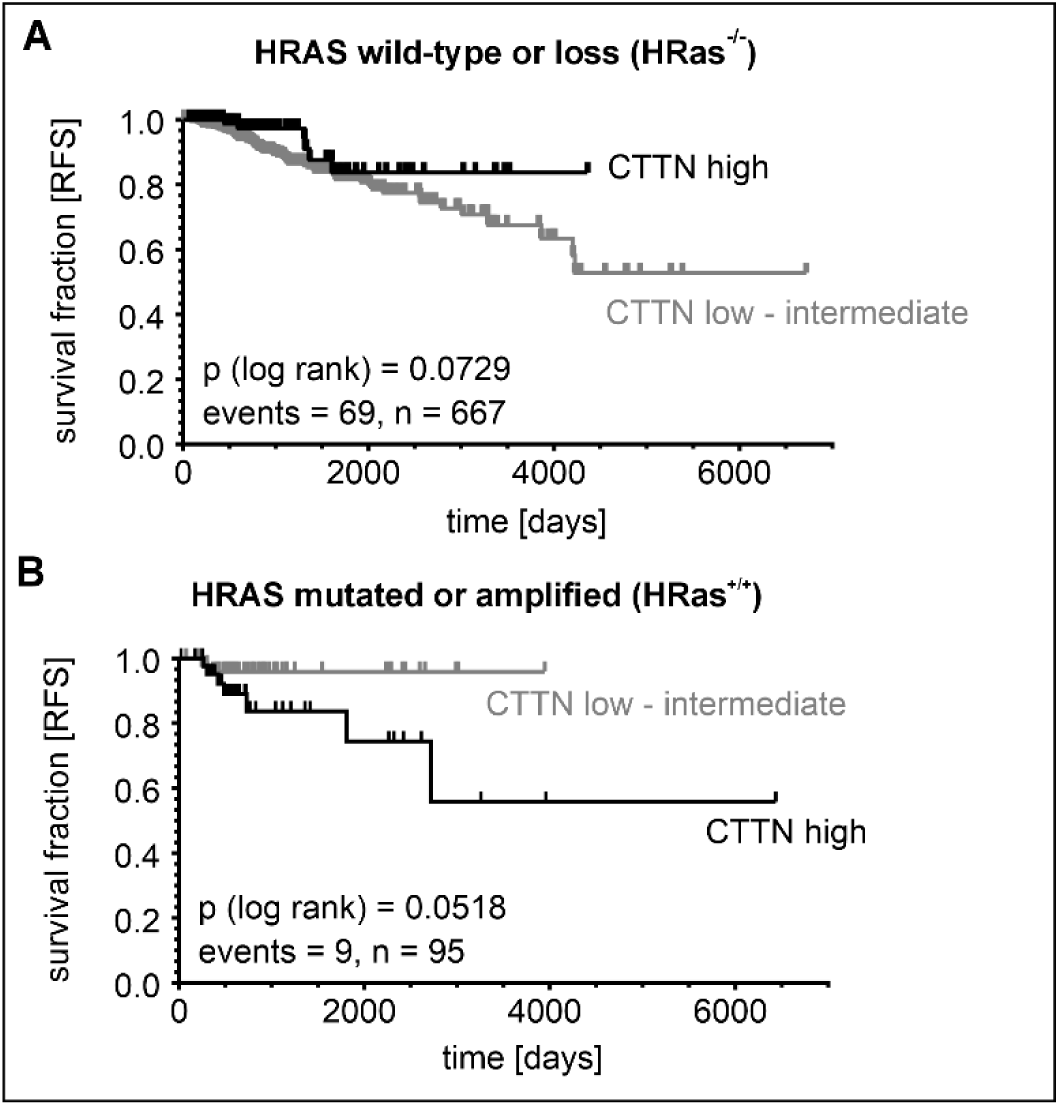
High cortactin and HRas gene expression predict poor survival of breast cancer patients. Univariate Kaplan-Meier curves of recurrence-free survival (RFS) of breast cancer patients stratified by HRas status in wild-type / loss: HRas^-/-^ (A) or mutated / amplified: HRas^+/+^ (B) and cortactin (CTTN) mRNA expression (low or high).

## Discussion

This study elucidated the poorly understood downstream signaling cascades that drive cellular programs of HRas-induced breast cancer invasion. It is characterized by locally restricted breakdown of the BM barrier, followed by cell transmigration into the TME. Our findings underline the remarkable context specificity of Ras GTPases, which act as critical molecular switches for cell fate decisions. Thereby, different Ras family members orchestrate diverse signaling pathways that control homeostatic cell existence but also malignant progression ^29, 56^. However, the underlying molecular mechanisms by which HRas coordinates the transition of cells from *in situ* to invasive breast carcinomas remained unclear.

To address early steps of metastatic breast cancer progression, we used MCF10A-based breast cell spheroids as simplified models of breast gland microtissue architecture. Without oncogenic HRas, these spheroids are well-established and appreciated for studying both normal breast gland morphogenesis and tumorigenic cell transition ^57^. A significant advantage of MCF10A spheroids is their ability to build and maintain basoapically polarized architectures ^58^. MCF10A spheroids endogenously produce and actively assemble a basement membrane (BM) scaffold that forms a physiologically meaningful mechanical force barrier ^8^. Previous work further demonstrated that invasive breast cells stress and disrupt the BM barrier, physically and proteolytically, to transmigrate into the surrounding TME ^5, 14^.

Since our spheroid model lacks any accessory myoepithelial cells or TME-associated stromal cells, our study strictly focused on elucidating bidirectional cell-ECM signal processing driven by oncogenic HRas signaling. We used MCF10A-based breast spheroids with inducible HRas downstream signaling ^33^. This 3D spheroid model resembled very early stages of oncogenic HRas transformation and revealed how morphologically normal, yet Ras hyperactivated epithelial cells initiate invasive behavior in response to matrix stiffening.

Tumor-associated ECM stiffening is a key driver of epithelial invasion, including in non-transformed breast spheroids ^5, 59, 60^. Remarkably, we found that oncogenic HRas activation partially bypasses this tumor-specific ECM cue, inducing effective invasive transitions in physiologically compliant, hence, healthy-like environment (cf. Fig. 1D). This finding contrasts with the inherently tumor-suppressive effects of normal matrix compliance on breast epithelial cells ^5, 14, 61^. However, our finding is consistent with prior reports of HRas-driven invasion under compliant conditions ^62^, highlighting its capacity to override invasion-protective mechanotransduction cues. Conversely, tumor-like ECM stiffness strongly amplified HRas-driven invasion and accelerated its onset (cf. Fig. 2D).

Previous studies showed that Ras hyperactivation during early morphogenesis induces EMT over several days in poorly polarized spheroids ^62, 63^. In contrast, HRas activation in fully polarized spheroids triggered a rapid invasive transition within hours. To our knowledge, such rapid HRas-driven invasion dynamics have not been previously reported. These findings highlight the potent effect of temporally controlled HRas activation in differentiated, non-tumorigenic epithelium. In vivo, ECM stiffening occurs not only during malignant progression but also in benign fibrocystic remodeling ^64^. These findings raise the possibility that oncogenic HRas activity may promote early invasion in otherwise histologically unsuspicious breast tissue ^65–67^.

At the cellular level, our findings indicated altered mechanotransductive responses associated with HRas activation. HRas^on^ cells developed actin-rich microspikes (MS) at the BM-ECM interface (cf. Fig. 3A), consistent with their described role as dynamic mechanosensory units in MCF10A spheroids: without oncogenic HRas signaling, MS have been shown to convert into contractile stress fibers (SF) that associate with an EMT-like phenotype, increased physical BM-stress and disruption ^23^. In contrast, HRas^on^ cells retained MS morphology without forming SFs. This new finding suggests the engagement of an alternative, SF-independent mechanotransductive pathway. However, the exact mechanism of HRas-modulated ECM-sensing remained elusive and needs further in-depth investigation.

These observations prompted us to further dissect the cellular strategies underlying HRas-mediated invasion: SF-lack coincided with the absence of increased non-muscle myosin II-driven cell contractility – a cellular force mode typically linked to mechanical BM stress, disruption and cell invasion ^23, 68^. Although such forces are usually amplified in response to pathological ECM stiffening ^14, 38^, TFM revealed only marginal contractile activity in HRas^on^ spheroids even on tumor stiffness, insufficient to explain their high invasiveness. This surprising finding highlighted the limited role of SF-mediated contractility and bulk traction forces in HRas-induced BM invasion and suggests the involvement of a non-canonical, non-contractile BM breaching mechanism.

In addition to actomyosin contractility, matrix metalloproteinase (MMP)–mediated BM degradation hallmarks epithelial invasion ^69, 70^. However, pharmacological inhibition of key pro-invasive MMPs did not impair HRas-driven invasion ^39, 69^ and expression of the cell membrane-anchored MT1-MMP (MMP14) remained unchanged upon HRas activation. This contrasts with the MMP dependency observed under oncogenic EGFR signaling in the MCF10 wild-type spheroid model ^5^. Recent work reported that oncogenic Ras induces steady gradual epithelial disintegration via hepsin-mediated laminin-332 degradation in soft matrix environments ^71^. The highly effective invasion onset of HRas^on^ spheroids on stiff substrates suggested that serine proteases like hepsin could have contributed but are not the primary mechanism. Our data demonstrated the contextual activation of distinct HRas invasion programs.

In search of the still unknown invasion mechanism, we examined the CBS-interface and discovered prominent cortical triplets (CTs) in HRas^on^ cells that most likely started the invasive transition. These actin-enriched patches were hallmarks of invasion and characterized by reinforced cortical actin, high cortactin accumulation, and localized BM disruption. Consistent with cortactin incorporation at the basolateral cell cortex, *CTTN* gene expression was substantially upregulated by HRas signaling. Cortactin is known to promote breast cancer cell migration ^72^, and invadopodia formation ^9, 73^. Typical invadopodia are finger-shaped actin-rich cell protrusions with MMP14-dependent proteolytic activity that widen collagen pores to facilitate invasion ^10, 74^. However, CT-positive HRas^on^ cells did not form such structures, and invasion proceeded independent of MMP activity. Instead, HRas^on^ cells assembled CT patches at basolateral cell surfaces.

Our data supported an HRas-driven invasion mechanism in which reinforced cortical actin provides mechanical input for BM penetration, independent of MMP proteolysis and invadopodia activity. Reinforced cortical actin coincided with dynamic pushing and pulling motions, spheroid compaction, and progressive BM densification (cf. Fig. 4), shape changes indicative of mechanical stress accumulation at the BM. While previous work demonstrated that invasive breast spheroids can exert substantial BM-stressing actomyosin II forces ^5^, here TFM revealed only low-level contractile stresses with minimal substrate deformations (strain energy ∼10 fJ) across all conditions and independent of HRas status. These values fell within the range observed in non-invasive breast spheroids (1–32 fJ) (7) and were therefore insufficient to account for the pronounced invasive phenotype. This suggests that HRas-driven invasion relies on a distinct mode of force generation. Consistently, focal disruption of reinforced cortical actin structures using laser-assisted nanosurgery demonstrated locally elevated cortical tension at these sites (cf. Fig. 6H). Based on the curvature-dependent coupling between cortical tension and outward pressure ^75^ this elevated tension is expected to generate localized outward-directed forces at the CBS interface. Notably, the contractile forces generated by the actin cortex are balanced by the BM. Therefore, little or no lateral stress is transmitted to the substrate which is in line with our TFM results.

We identified Arp2/3-mediated actin polymerization and myosin I activity as key drivers of localized pro-invasive forces. Mechanistically, HRas engages Src kinase to promote CT formation. Src kinase, through phosphorylation of cortactin at Tyr421, likely links HRas signaling to Arp2/3-dependent cytoskeletal remodeling at sites of BM disruption, where cortactin co-localizes with reinforced cortical actin ^49, 53, 76^.

Arp2/3-driven actin branching promotes filament network densification, mechanical reinforcement ^77^, and enhanced protrusive capacity of migrating cells ^78^ and has been shown to mediate BM disruption independently of MMPs or actomyosin II in *C. elegans* embryogenesis ^79^. Recent work further demonstrated that myosin I synergizes with Arp2/3 to amplify force generation in branched actin networks, likely by reorganizing filament architecture into a mechanically more efficient configuration ^80^. Consistently, inhibition of either pathway reduced invasion of HRas^on^ cells to levels comparable to HRas^off^ controls, identifying Src, Arp2/3, and myosin I as key effectors of HRas-driven BM disruption and potential therapeutic targets. Whether elevated cortical tension and actin polymerization-driven force generation act interdependently or represent parallel mechanical inputs to BM disruption remains to be determined.

Together, these data reveal a coordinated downstream signaling cascade driving BM disruption: (1) oncogenic HRas activates Src kinase, which (2) phosphorylates the actin binding protein cortactin at Tyr421, leading to (3) Arp2/3 stabilization, (4) enhanced cortical actin branching and polymerization forces. Ultimately, (5) these myosin I-synergized forces foster BM disruption and invasive transition. Although additional candidates such as ephrin-B4 and aurora kinase A were identified by kinase profiling, their inhibition had no functional outcome, underscoring the specificity of the HRas–Src–cortactin–Arp2/3-myosin I axis for cell invasion. These findings highlight that oncogenic Ras signaling integrates diverse pathways with context-dependent cellular consequences ^81, 82^.

To evaluate the translational relevance of our mechanistic findings, we assessed clinical breast cancer datasets. Given the well-established context specificity of HRas signaling, it was critical to determine whether this invasion-related pathway also manifests in patient tumors and correlates with clinical outcome. HRas overexpression is recognized as an independent marker of poor prognosis ^83^ and activating HRas mutations are associated with adverse survival in breast cancer patients. We therefore stratified tumor cases based on HRas copy number and mutation status and analyzed co-expression patterns with cortactin, a marker of metastatic potential at both the transcript and protein levels as well ^84, 85^. Although the low prevalence of HRas mutations in breast cancer ^2^ limited the size of available patient cohorts, our *in silico* analysis revealed a significant association between high co-expression of HRas-cortactin and reduced recurrence-free survival (RFS), which fits our *in vitro* observation of HRas-induced cortactin transcription. Conversely, tumors with high cortactin but low HRas expression indicated improved RFS. This suggests a directionally dependent functional interaction between these two effectors in tumor progression. Further supporting evidence is that the amplification of ARP2/3 complex subunits was associated with poor prognosis in the HRas-mutated patient subgroup. Among the seven ARP2/3 subunits ^55, 86^, two structural core elements (ARPC2 and ARPC4) ^87^ and one key mediator of actin nucleation and assembling (ARPC1B) ^88, 89^, each correlated significantly with reduced recurrence-free survival. These findings strengthen the link between the HRas–cortactin–Arp2/3 signaling axis and breast cancer progression, supporting its relevance for patient outcome.

### Limitations of the study

While 3D spheroid models enabled functional dissection of the HRas–Src–cortactin–Arp2/3 axis, our study is inherently based on a cell culture model, whose simplified architecture cannot fully recapitulate the complexity of HRas-mutated or -hyperactivated tumors *in vivo*. This reflects a fundamental technical limitation, as precise spatiotemporal control of HRas activity-required to dissect early oncogenic events- cannot currently be achieved *in vivo*: Importantly, breast cell spheroids enabled causal dissection of early oncogenic processes and their associated cytoskeletal dynamics at a resolution not experimentally accessible using for instance rodent *models*. Nevertheless, the absence of accessory myoepithelial cells and other TME-associated stromal components precludes to resemble reciprocal epithelial–stromal feedback loops that may influence epithelial cell behavior ^90^. Future studies incorporating such heterotypic interactions will be important to further refine our understanding of HRas-driven breast cancer progression. Accordingly, validation in complementary *in vivo* systems, such as inducible HRas^on/off^ rodent xenografts, will be an important step to extend and validate these mechanistic and prognostic insights.

## Concluding remarks

Our findings reveal that oncogenic HRas signaling rewires mechanosensing of tumor microenvironmental stiffness, cell mechanics and actin forces to drive BM disruption and early cell invasion in phenotypical non-malignant breast epithelial cells. We show that this early-stage invasive transition is mechanistically dependent on Src kinase phosphorylated Tyr421 cortactin, Arp2/3, actin polymerization and myosin I forces. Intriguingly, these cellular switches could represent promising therapeutic targets to counteract HRas-driven invasion at early stages. While BM-covered breast spheroids simplify *in vivo* complexity, the correlation of HRas downstream effector expression with reduced recurrence-free survival in breast cancer cohorts supports the clinical relevance of this newly discovered oncogenic signaling pathway (see Fig. 8). Finally, these results deepen our understanding of how spontaneous oncogene activation perturbs mechanobiological homeostasis during early breast cancer progression.

**Figure 8:**
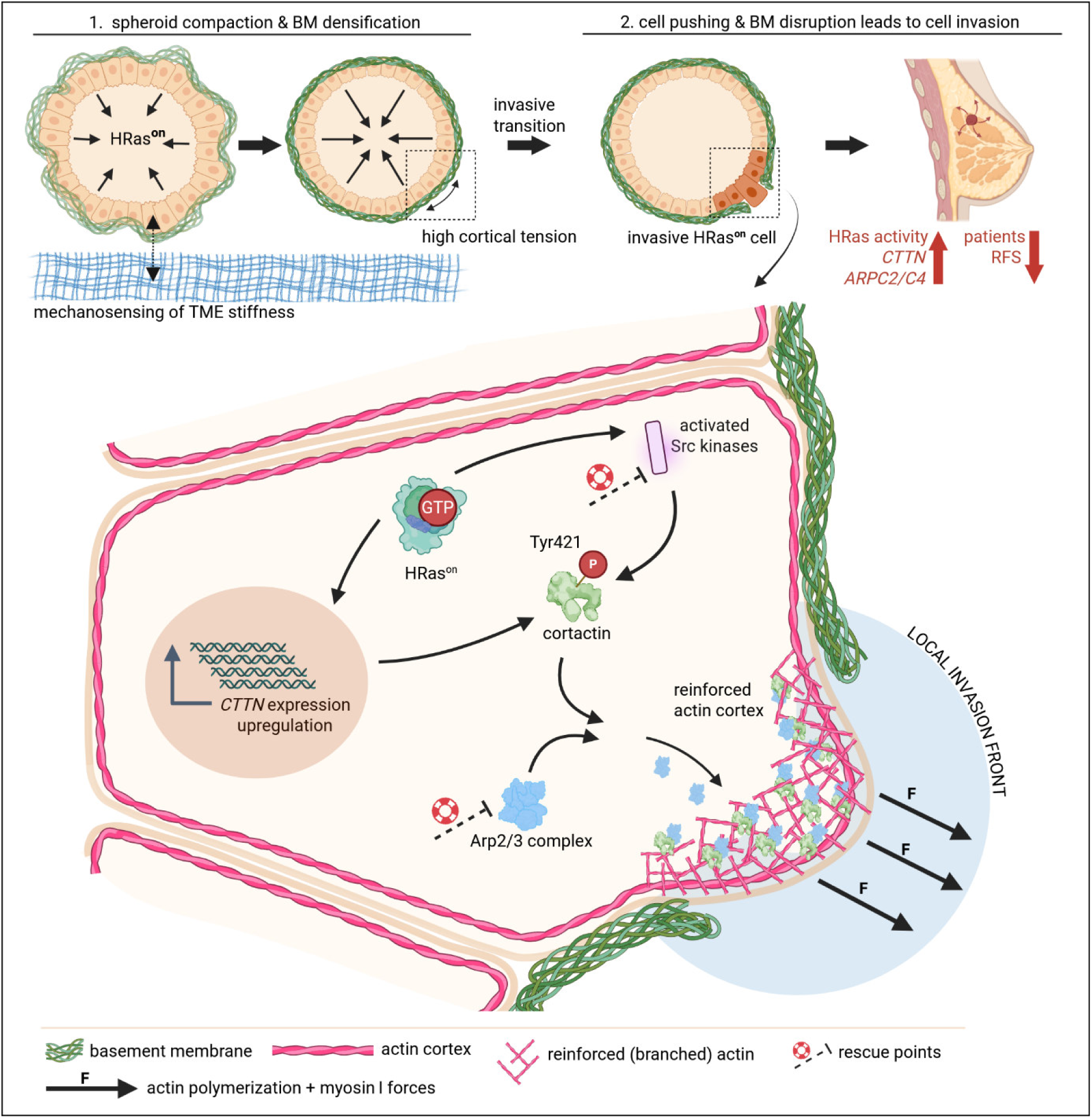
Oncogenic HRas rewires myosin I and actin polymerization forces to drive BM invasion. (1) Oncogenic HRas activation in breast epithelial cells reprograms epithelial responses to TME stiffening, resulting in cell compaction, BM densification and elevated cortical tension. (2) HRas activated cells initiate invasion via a Src–cortactin signaling axis: Src phosphorylates cortactin at Tyr421, which recruits and stabilizes the Arp2/3 complex at the basolateral cortex to establish a reinforced actin-rich invasion front. Arp2/3-dependent actin polymerization together with myosin I-generated forces deform and ruptures the BM barrier, enabling cell transmigration. HRas-dependent upregulation of cortactin expression fosters this invasion program. Pharmacological inhibition of Src, Arp2/3, or myosin I restore a non-invasive phenotype. Clinically, elevated expression of HRas effectors (cortactin and ARP2/3 subunits) correlates with reduced recurrence-free survival in HRas-amplified or -mutated breast cancers. Schematics are not to scale. Created with BioRender.com

## Materials and Methods

### Cell Maintenance

MCF10A wildtype cells (purchased from ATCC), MCF10A_ER:HRas^G12V^ and MCF10A_HRas^G12V^ cells described in ^34^ and kindly provided by Buzz Baum ^35^ were maintained in culture dishes under standard culture conditions (37 °C, 5% CO_2_) in DMEM/F12 growth medium (ThermoFisher Scientific) containing 5% horse serum (ThermoFisher Scientific) or steroid hormone free horse serum (c.c.pro) (in all experiments from Fig. 3 onwards), 0.5 µg/mL hydrocortisone, 100 ng/mL cholera toxin, 20 ng/mL EGF, 10 µg/mL insulin (all Sigma Aldrich), 100 U/mL penicillin and 100 µg/mL streptomycin (both ThermoFisher Scientific). For spheroid morphogenesis, assay media with adapted compositions were used (see below).

### Spheroid morphogenesis and isolation from EHS matrix

Single cells were seeded on top of growth factor reduced EHS matrix (Geltrex, ThermoFisher Scientific) and cultured for ten days as described in ^23^. MCF10A wildtype cells were cultivated from day one to day nine and MCF10A cells with HRas constructs from day one to day six in assay medium with 5 ng/mL EGF and 1% horse serum. Assay medium without EGF was used for wildtype cells from day nine to day ten and for MCF10A cells with HRas constructs from day six to day ten. For further analyses, spheroids were isolated from EHS matrix and washed with ice-cold PBS and incubated in 2 mL ice-cold cell recovery solution (CRS) (BD Biosciences) for 30 min (4 °C) to depolymerize the EHS matrix. For inhibition experiments, inhibitors were added 30 min prior to isolation. Next, spheroids were washed with fresh EGF-free assay medium, picked under a stereo microscope and seeded onto 35 mm cell culture dishes, either with glass bottoms or glass bottoms with a layer of silicone elastomer (see below). Spheroids adhered to these EHS-coated substrates for 15 min (37 °C, 5% CO_2_) and were subsequently covered with 2 mL EGF-free assay medium. The time point of adding media was defined as assay start.

### Preparation of elastomeric substrates and functionalization with EHS coating

Spheroids were transferred on an 80 µm thick layer of cross-linked PDMS silicone elastomer substrate (Sylgard 184, Dow Corning) with a Young’s modulus of 16 kPa. Preparation of these substrates was done as described in ^91^. In brief, layer thickness was set by spin coating on 100 µm thin cover slips (Cover Slip, Ø22 mm, #0, Menzel-Gläser). Silicone-coated cover slips were glued to the bottom of 3.5 cm Petri dishes to cover predrilled 1.8 cm holes and cross linked for 16 hours at 60 °C. Young’s modulus silicone elastomers was determined as described previously ^92^. For TFM, fluorescent beads (FluoSpheres, carboxylate-modified, 0.2 µm, red, ThermoFisher Scientific) were immobilized on top of elastomeric substrates, as described elsewhere ^93^. Before spheroid transfer, elastomeric substrates and glass substrates (Cover Slip, 24 x 24 mm, HP, Menzel-Gläser) were functionalized for adhesion with 600 µL of non-gelling EHS protein solution (20 µg/mL) in ice-cold PBS for 18 hours at 4 °C.

### Biochemical treatments

Spheroids were transferred on elastomeric substrates and incubated with EGF-free assay medium after 15 min. For activation of HRas, a final concentration of 1 µM OHT (H7904, Sigma-Aldrich; dissolved in EtOH) was used (final EtOH concentration = 0.002%). For inhibition of cellular myosin II activity, a final concentration of 25 µM blebbistatin (B0560, Sigma-Aldrich; dissolved in DMSO) was used (final DMSO concentration = 0.29%). For inhibition of matrix metalloproteinases, a final concentration of 20 µM marimastat (M2699, Sigma-Aldrich; dissolved in DMSO) was used (final DMSO concentration = 0.03%). For inhibition of ephrin type-B receptor 4, a final concentration of 5 µM NVP-BHG712 (HY-13258A, MedChemExpress; dissolved in DMSO) was used (final DMSO concentration = 0.01%). For inhibition of aurora kinase A, a final concentration of 5 µM MK-5108 (HY-13252, MedChemExpress; dissolved in DMSO) was used (final DMSO concentration = 0.025%). For inhibition of myosin I, a final concentration of 20 µM Pentachloropseudilin (HY-115669, MedChemExpress; dissolved in DMSO) was used (final DMSO concentration = 0.007%). For inhibition of Src kinase, a final concentration of 5 µM PP2 (1407, Tocris; dissolved in EtOH) was used (final EtOH concentration = 0.05%). For inhibition of Arp2/3 complex, a final concentration of 40 µM CK-666 (3950, Tocris; dissolved in EtOH) was used (final EtOH concentration = 0.04%).

### Immunofluorescent staining

Spheroids were fixed for 20 min with 3.7% paraformaldehyde in cytoskeleton-buffer (CB: 5 mM EGTA, 5 mM glucose, 10 mM MES, 5 mM MgCl_2_, 150 mM NaCl, 1 g/L streptomycin; all Sigma-Aldrich), washed for 5 min with 20 mM glycine in CB (Sigma-Aldrich), and permeabilized with 1% Triton-X 100 in CB (Sigma-Aldrich) at RT. After one washing step with CB, non-specific antibody binding was blocked with 5% skim milk powder (Sigma-Aldrich) and 1% AffiniPure F(ab’)_2_ fragment goat anti-mouse IgG (115-006-006, Jackson ImmunoResearch, West Grove, PA, USA) in CB for 2 hours at RT. Spheroids were incubated overnight at 4 °C with primary antibodies anti-collagen IV (1:500, abcam, ab6586), anti-GM130 (1:500, BD Biosciences, 610822, clone 35), Alexa Fluor 488-conjugated anti-laminin-5 antibody (1:1000, Sigma-Aldrich, MAB19562X; clone D4B5), anti-cortactin (p80/85) (1:300, Sigma-Aldrich, 05-180-I, clone 4F11), anti-phospho-Tyr421-cortactin (1:300, ThermoFisher, 44-854G), anti-pERK (Phospho-p44/42, Thr202/Tyr204) (1:200, Cell Signaling, 9101S), anti-MT1-MMP (1:200, Abnova, MAB12762, clone 133CT15.10.5.1) diluted in 1% skim milk powder in CB followed by three washing steps with CB.

Conjugated secondary antibodies chicken anti-rabbit IgG Alexa Fluor™ 488 (ThermoFisher Scientific, A21441), goat anti-rabbit IgG Alexa Fluor™ 488 (ThermoFisher Scientific, A11008), goat anti-rabbit IgG Alexa Fluor™ 488 (ThermoFisher Scientific, A31556), donkey anti-rabbit IgG Alexa Fluor™ 546 (ThermoFisher Scientific, A10040), donkey anti-mouse IgG Alexa Fluor™ 546 (ThermoFisher Scientific, A10036), were diluted (1:1000) in 1% skim milk powder in CB and incubated for 1 h at RT followed by two washing steps with CB. Phalloidin-Atto 633 (Sigma-Aldrich, 68825) or 488 (Sigma-Aldrich, 49409) labeling was done parallel to secondary antibody administration (1:1000). Nuclei were stained with 1:1000 DAPI in CB (ThermoFisher Scientific, NucBlue™ Fixed Cell ReadyProbes™) or 1:1000 DRAQ5™ in CB (ThermoFisher Scientific, 62251) for 10 min at RT. To enable reliable signal quantification, fixation, staining, and imaging were performed in an identical manner across experiments, maintaining sequence and procedures for each step. For live cell imaging, spheroids were stained in assay medium with a final concentration of 10 nM SiR-actin (Spirochrome, SC001, solved in DMSO, final DMSO concentration = 1×10^-5^%) for 2.5 hours, isolated from EHS matrix and incubated with 1:100 Alexa Fluor 488-conjugated anti-laminin-5 antibody (Sigma-Aldrich, MAB19562X; clone D4B5) for 1 hour at 4 °C. Spheroids were then transferred onto silicone elastomer substrates.

### Confocal microscopy and image processing

Fixed and stained spheroids were imaged at room temperature, and live-cell imaging was performed at 37 °C and 5% CO₂ using an inverted confocal laser scanning microscope (LSM880 with Airyscan detector; Carl Zeiss). The system was equipped with a 40× LD C-Apochromat water immersion objective (NA 1.1; Zeiss) and standard laser lines (405, 488, 561, and 633 nm) with appropriate filter sets. Images were acquired using (Fast) Airyscan detection and processed by Airyscan processing in ZEN 2.3 Black (Carl Zeiss). Three-dimensional visualizations were generated using PyVista (93) and Imaris 10.2 (Oxford Instruments).

### Laser-assisted nanosurgery of cortical actin structures

Scanner-based laser ablation was performed using a nanosecond-pulsed 355 nm laser (UGA-42 Firefly/Caliburn DPSL-355/42/CLS; Rapp OptoElectronic) coupled to the LSM described above. A mean output power of 8.4 mW was used. A 20x air objective (NA 0.8, Carl Zeiss) was used for all ablation experiments. Spheroids were stained with live actin dye SiR-actin (see section Immunocytochemistry) for 3 hours (parallel to OHT treatment). Visibly thickened cortical regions were used as samples for reinforced actin sites while thin regions served as control, non-reinforced sites. Ablation was performed using an unfilled rectangular region of interest (42 x 3 pixels; 8.11 x 0.58 µm at the given image scale) oriented perpendicular to the actin cortex (step size 5, 10 repetitions; sequence mode SysCon, Rapp OptoElectronic). Object illumination and therefore ablation took 187 ms. Images were acquired every 200 ms in Fast-Airyscan mode. Imaging started ≥1.5 s prior to ablation for baseline calculation (see Supplemental Information).

### Live-cell imaging for invasion assay and TFM

Experiments were carried out at 37 °C and 5% CO_2_ using an inverted microscope (Axio Observer, Zeiss), equipped with an Axiocam 712 mono camera (Zeiss) and an EC Plan-Neofluar 40x oil immersion objective (PH3, NA 1.3, Zeiss). Cells were imaged in phase-contrast and fluorescent beads with a LED module (Colibri 7, Zeiss) at 548 nm and excitation and emission filter settings of 538 – 562 nm and 570 – 640 nm, respectively. Time increment of image acquisition was 20 min (image pixel size = 0.17 µm). Tangential substrate deformations were visualized by tracking fluorescent beads to calculate strain energy, the elastic energy of the cell-deformed substrate (see Supplemental Information).

### Digital image processing of confocal laser scanning microscopy micrographs

All routines for analyses were developed in Python 3.12, including quantification of pERK, MT1-MMP, BM and CT signal, of compactness and of actin after laser ablation, and are described in detail in Supplemental Methods.

### Kinome profiling analysis

Kinase profiles were determined using the PamChip^®^ peptide tyrosine kinase (PTK) microarray system or the PamChip^®^ Ser/Thr Kinase assay (STK) on PamStation^®^12 (PamGene International, s-Hertogenbosch, The Netherlands). Each PTK-PamChip^®^ array contained 196 individual phospho-sites(s) that are peptide sequences derived from substrates for tyrosine kinases. Each STK-PamChip^®^ array contained 144 individual phospho-site(s) that are peptide sequences derived from substrates for Ser/Thr kinases. Each peptide on the PAMChip is a 15-amino acid sequence representing a putative endogenous phosphorylation site, which functions as a kinase substrate. Sample preparation and phosphorylation detection are described in detail in the Supplemental Methods section.

### RT-qPCR analysis

Spheroids on 16 kPa elastomeric substrates, incubated 270 min with EtOH or OHT, were mechanically pipetted in solution with assay medium (without EGF). Total RNA from spheroids was isolated through column chromatography according to the manufacturer’s protocol (QIAGEN, miRNeasy Tissue/Cells Advanced Mini Kit, 217604) and quantified using a NanoDrop spectrophotometer (ThermoFisher). Total RNA samples were reverse transcribed using RT2 First Strand Kit (QIAGEN, 330401) according to the manufacturer’s protocol. Generated cDNA was used for qRT-PCR array with cytoskeleton regulators (QIAGEN, PAHS-088Z). PCR measurements were conducted on a QIAquant 96 station (QIAGEN) and analyzed according to the array manufacturer’s instructions and their *GeneGlobe* online tool. A C_T_ cut-off of 35 was used and normalization was done with two genes based on minimal difference of geometric means across samples (<1 C_T_).

### TCGA data and survival statistics

Publicly available breast cancer data from The Cancer Genome Atlas (TCGA) network (Breast Invasive Carcinoma (Firehose Legacy) (1,108 samples)) were used to determine the clinical impact of CTTN and ARP expression in context of genetic HRAS alterations comprising transcriptomic (RNASeqV2 data: *CTTN, ACTR2, ACTR3, ARPC1A, ARPC1B, ARPC2, ARPC3, ARPC4, ARPC5, ARPC5L*), genomic (HRAS mutation status and HRAS-associated copy number variation) and clinical follow-up data of overall n=1,082 primary breast cancer samples (https://gdac.broadinstitute.org/runs/stddata__2016_01_28/data/BRCA/20160128/). Data was accessed by using the cBio Cancer Genomics Portal (http://cbioportal.org) ^94^ and analyzed by using SPSS software version 29.0.0.0 (SPSS Inc., Chicago, USA). Survival curves for recurrence free survival (RFS) were stratified by genetic HRAS alterations (copy number variation and mutations), calculated using the Kaplan-Meier method with log-rank. RFS was measured from surgery until relapse (local/distant) and was censored for patients without evidence of tumor recurrence at the last follow-up date.

### Figure preparation

Figure 8 was created in BioRender. Platz-baudin, E. (2026) https://BioRender.com/pl9dmm8

## Resource availability

## Lead contact

Requests for further information and resources should be directed to and will be fulfilled by the lead contact, Erik Noetzel.

## Materials availability

All unique/stable reagents generated in this study are available from the lead contact without restriction.

## Data and code availability

All data needed to evaluate the conclusions in the paper are present in the paper and/or the supplemental information. Any additional information required to reanalyze the data reported in this paper is available from the lead contact upon request, Erik Noetzel.

### Acknowledgments

We thank Helen Matthews (Sir Henry Dale Fellow School of Biosciences, University of Sheffield UK) and Buzz Baum (MRC Laboratory of Molecular Biology, Cambridge UK) for kindly providing the HRas cells and Frederik Rastfeld (IBI-2 Research Center Jülich) for his practical support. This work was supported by the Corona Foundation (S199/10084/2021), and by the Deutsche Forschungsgemeinschaft (DFG) (SFB TRR219 – Project-ID 322900939; subproject M07 and INST 222/1598-1 FUGG; Projekt: 566187929) to E.P.C.v.d.V. Funded by the Deutsche Forschungsgemeinschaft (DFG, German Research Foundation) 363055819/GRK2415 to EN.

## Author contributions

Conceptualization, EN, EPB; methodology, EPB, JE, YH, GD, RW, MR, RM; formal analysis, EPB, GD, RW, EPCvdV, MR; funding acquisition, EN; project administration, EN, RM; investigation, EPB, JE, YH, EPCvdV, MR, RW; supervision, EN, RM; validation, EPB; visualization, EPB, EN, EPCvdV, MR; writing – original draft, EPB, EN, MR; writing – review & editing, EPB, RM, EN, MR.

## Declaration of generative AI and AI-assisted technologies in the writing process

During the preparation of this work the authors used ChatGPT5.2 to check grammar, spelling and clarification of phrases in parts of the manuscript. After using this tool, the authors reviewed and edited the content as needed and take full responsibility for the content of the published article.

## Supplemental Information

Supplemental Video S1: HRas^on^ spheroid in invasion assay on 16 kPa. Phase contrast image series of an HRas-activated spheroid. After 12 hours the first cell invades the surrounding. Corresponds to Figure 2C.

Supplemental Video S2: BM breach of HRas^on^ spheroid on 16 kPa. Series of confocal images (F-actin in magenta and BM/laminin-332 in cyan) with a time increment of 15 min. Cells of the spheroid retracted during the initial 4 hours, followed by a pushing phase that culminated in BM disruption at 6 hours. Corresponds to Figure 4A.

Supplemental Video S3: Laser ablation of a non-reinforced cell cortex (control sites) at an HRas^on^ spheroid on 16 kPa. Series of confocal images (F-actin in grey) with a time increment of 200 ms. In the video, ablation started after relative setpoint t = 0 ms. Corresponds to Figure 6E.

Supplemental Video S4: Laser ablation of a reinforced cell cortex at an HRas^on^ spheroid on 16 kPa. Series of confocal images (F-actin in grey) with a time increment of 200 ms. In the video, ablation started after relative setpoint t = 0 ms. Corresponds to Figure 6F.

## Supplemental Methods

### Traction force microscopy

Maps of cell-induced traction stresses were calculated by regularized least-square fitting to the mechanical response of an elastic layer of 80 µm thickness on rigid substrates ^95^. From these maps, strain energy was calculated as a scalar measure of overall mechanical activity. These calculations were done as described in previous work ^95, 96^ and integrated over an area of 38,007 µm^2^ to cover the entire region of influence of a spheroid. The required algorithms were implemented in MatLab (R2015a, The MathWorks Inc.). Spheroids were analyzed for at least 24 hours. All displacements (strains) and forces (stresses) were calculated with reference to the first image of the series (t = 0 hours). Changes in strain energy were therefore determined with respect to that reference state. For baseline correction, calculated strain energy values were subtracted by the mean of 3 time points of at least two cell-free positions per experiment (see Supplementary Fig. S2).

### Python routines for image processing

**For quantification of pERK signals**, images of pERK staining with nuclei and F-actin co-staining (pixel size=0.09 µm) were processed using a Gaussian filter (σ=1). To extract the outer spheroid shape, a maximum intensity projection (MIP) through the three channels was generated that used the highest intensity value at each pixel. Using the threshold selection method developed by Otsu ^97^ and scaling of the resulting threshold by a factor of 0.5, a binary spheroid mask was generated of the MIP image. The resulting mask was refined using a morphological closing operation (cross-shaped structuring element of 3×3 pixels, applied iteratively 50 times), hole filling to eliminate internal gaps, a morphological opening operation (cross-shaped structuring element of 3×3 pixels, applied iteratively 20 times) and selection of the largest connected component.

To identify individual cell masks within the spheroid, the nuclei channel was segmented using the Otsu thresholding method (scaled by a factor of 0.5). Morphological operations were then applied to clean the segmentation: opening with a square-shaped structuring element (5 pixels in size), removal of objects smaller than 2500 pixels, closing with the same rectangular structuring element and hole filling. A watershed algorithm was applied to all objects, to separate touching or clustered nuclei. This involved computing the distance transform of the binary nuclei mask and identifying local minima as markers for the watershed. The boundaries generated by the watershed segmentation were used to delineate individual cell regions. The mean intensity of pERK was calculated of individual cells inside the spheroid shape and averaged per spheroid.

**For quantification of MT1-MMP signals**, images of MT1-MMP staining were segmented using the threshold selection method developed by Otsu ^97^. The mean intensity was calculated of pixels above the threshold.

**For compactness measurements**, a maximum intensity projection (MIP) of z-planes (smoothed by a Gaussian filter; σ=3) was generated from the laminin-332 channel (pixel size=0.22 µm) for each spheroid. The minimum grayscale value of the MIP image was subtracted. Next, a threshold was computed using the triangle algorithm ^98^. This threshold was scaled by a factor of 1.2 and applied to separate signal from background. The resulting mask was refined using a morphological closing operation with a disk-shaped structuring element (15 pixels in radius), a morphological opening operation with a disk-shaped structuring element (5 pixels in radius), followed by hole filling to eliminate any internal gaps. If multiple objects remained after these steps, the largest connected component was selected and used as the final spheroid mask for calculation of compactness by

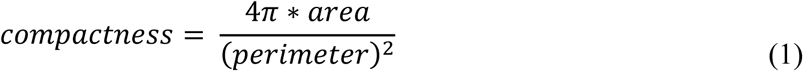

as the ratio of the measured area to the area of a circle with the same perimeter.

**For quantification of basement membrane signals**, individual z-planes were analyzed separately. The image of the laminin-332 channel (pixel size=0.11 µm) was smoothed using a Gaussian filter (σ=2). An Otsu threshold ^97^ was applied to separate signal from background. The resulting binary mask was refined using a binary opening operation with a disk-shaped structuring element (3 pixels in radius). For laminin-332 and collagen IV average intensities were calculated within the mask. Subsequently, intensities from different z-planes were averaged again.

**For analysis of cortical triplets (CT)**, individual z-planes were analyzed separately (pixel size=0.07 µm). The spheroid border was defined in the cortactin image channel as follows. Using the threshold selection method developed by Otsu ^97^, and scaling the resulting threshold by a factor of 0.5, background was separated from signal to generate a binary spheroid mask. The resulting mask was refined using a morphological closing operation (disk-shaped structuring element of 10 pixels in radius), hole filling to eliminate internal gaps, selecting largest connected component, a morphological opening operation (disk-shaped structuring element of 10 pixels in radius) and a morphological dilation operation (disk-shaped structuring element of 10 pixels in radius). For identification of localized, reinforced actin sites, the distance transform of the spheroid mask was calculated and only the outermost 54 pixels (equal to 4 µm at given image scale) were considered. In the following, this section is referred to as outer strip. Next, intensity thresholds were computed separately for each image plane based on the intensity distribution of the entire plane by

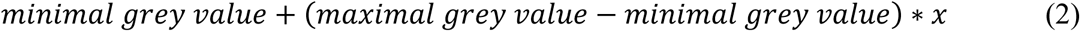

where x=0.25 for F-actin and cortactin channels and x=0.33 for collagen IV channel. In the F-actin channel (smoothed by a Gaussian filter; σ=5) bright objects (reinforced actin) were segmented by intensity thresholding (Eq. 2). Only objects within the outer strip were considered and objects smaller than 67 pixels were rejected. Next, we calculated the average intensity of cortactin within each identified F-actin object. If it exceeded the cortactin threshold (Eq. 2), this spot was identified as co-localized in F-actin and cortactin. Otherwise, it was rejected. In the final step, we searched for weakened BM at co-localized objects. Since the BM, and thus the collagen IV signal, surrounds the spheroid, the object was enlarged radially outwards by morphological dilation (size: 14 pixels, equal to 1 µm at given image scale). To do so, the center of mass of each object was determined in the previously calculated distance transform. Since the lowest values in the distance transform are located in the direction of the spheroid boundary, radial outwards extension was only performed for values that were below the center of mass value. Enlarged objects were rejected if the mean grey value of collagen IV was above the threshold. Conversely, the cortical triplet (CT) condition (F-actin and cortactin co-localization together with low BM signal) was fulfilled if the mean grey value of collagen IV was below the threshold. For each spheroid, CT frequency (CT positive pixels within outer strip to all pixels within outer strip) across all image planes was calculated.

For image visualizations, co-localization was calculated in Imaris (software version 10.2, Oxford Instruments) in “Threshold Selection Mode”. From the z-stack source image, one plane was randomly selected and thresholds were calculated by

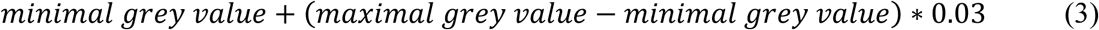

**For quantification of actin cortex retractions after laser-assisted rupture** (image pixel size = 0.19 µm), the cell edge contour was drawn manually in ImageJ (1.50b) and the resulting coordinates were replaced by a simplified polygonal path determined by an iterative point selection algorithm ^99^ with a tolerance of 2 px. From the remaining coordinate pairs, 20 equidistant reference points were interpolated along the smoothed path. As ablation was always performed in the image center but with angles adjusted to match the local cell edge orientation, ten rectangular-shaped sectors (polygon of four points; 3 px width and 15 px height) were arranged on each side of the center ablation sector in a row. In the next step, each segment was translated along its longitudinal axis such that the midpoint of the baseline contacted the approximated cell edge contour. The mean fluorescence intensity of each sector was calculated over time. The baseline per sector was calculated from the mean intensity values prior to ablation and set to 1.

### Sample preparation and phosphorylation detection for Kinome profiling analysis

Spheroids on 16 kPa elastomeric substrates, incubated 90 min with EtOH or OHT, were mechanically pipetted in solution with assay media and washed once in ice-cold PBS after respective treatments,, and lysed for 15 min on ice using M-PER Mammalian Extraction Buffer containing Halt Phosphatase Inhibitor and EDTA-free Halt Protease Inhibitor Cocktail (1:100 each; ThermoFischer Scientific). Three biological replicates were used per condition. Lysates were centrifuged for 15 min at 16,000x g at 4 °C in a pre-cooled centrifuge. Supernatants were aliquoted, snap-frozen in liquid nitrogen and stored at −80 °C. Protein quantification was performed with BCA Assay (ThermoFischer Scientific) according to the manufacturer’s instructions. For experiments, new aliquots were thawed.

**For the PTK assay**, 9.0 µg of protein was applied per array (n=3 per condition) and the assay was carried out using the standard protocol supplied by Pamgene International B.V. All reagents used for PTK activity profiling were supplied by Pamgene. Initially, to prepare the PTK Basic Mix, the freshly thawed supernatant was added to 4 µl of 10x protein PTK reaction buffer (PK), 0.4 µl of 100x bovine serum albumin (BSA), 0.4 µl of 1 M dithiothreitol (DTT) solution, 4 µl of 10x PTK additive, 4 µl of 4 mM ATP and 0.6 µl of monoclonal anti-phosphotyrosine FITC-conjugate detection antibody (clone PY20). Total volume of the PTK Basic Mix was adjusted to 40 µl by adding distilled water (H_2_O). Before loading the PTK Basic Mix on the array, a blocking step was performed applying 30 µl of 2% BSA to the middle of every array and washing with PTK solution for PamChip® preprocessing. Next, 40 µl of PTK Basic Mix were applied to each array of the PamChips®. Then, the microarray assays were run for 94 cycles. An image was recorded by a CCD camera PamStation^®^12 at kinetic read cycles 32–93 at 10, 50 and 200 ms and at end-level read cycle at 10, 20, 50, 100 and 200 ms.

**For the STK assay**, 2.0 µg of protein and 400 µM ATP were applied per array (n=3 per condition) together with an antibody mix to detect phosphorylated Ser/Thr. After incubation for an hour (30 °C) during which the sample was pumped back and forth through the porous material to maximize binding kinetics and minimize assay time, a second FITC-conjugated antibody is used to detect the phosphorylation signal. Imaging was done using a LED imaging system. Spot intensity at each time point was quantified (and corrected for local background) using the BioNavigator software version 6.3 (PamGene). Upstream Kinase Analysis (UKA) ^100^, a functional scoring method (PamGene) was used to rank kinases based on combined specificity scores (based on peptides linked to a specific kinase, derived from six databases) and sensitivity scores (based on treatment-control differences) (BioNavigator version 6.3).

## Supplemental Figures

**Supplementary Figure S1:**
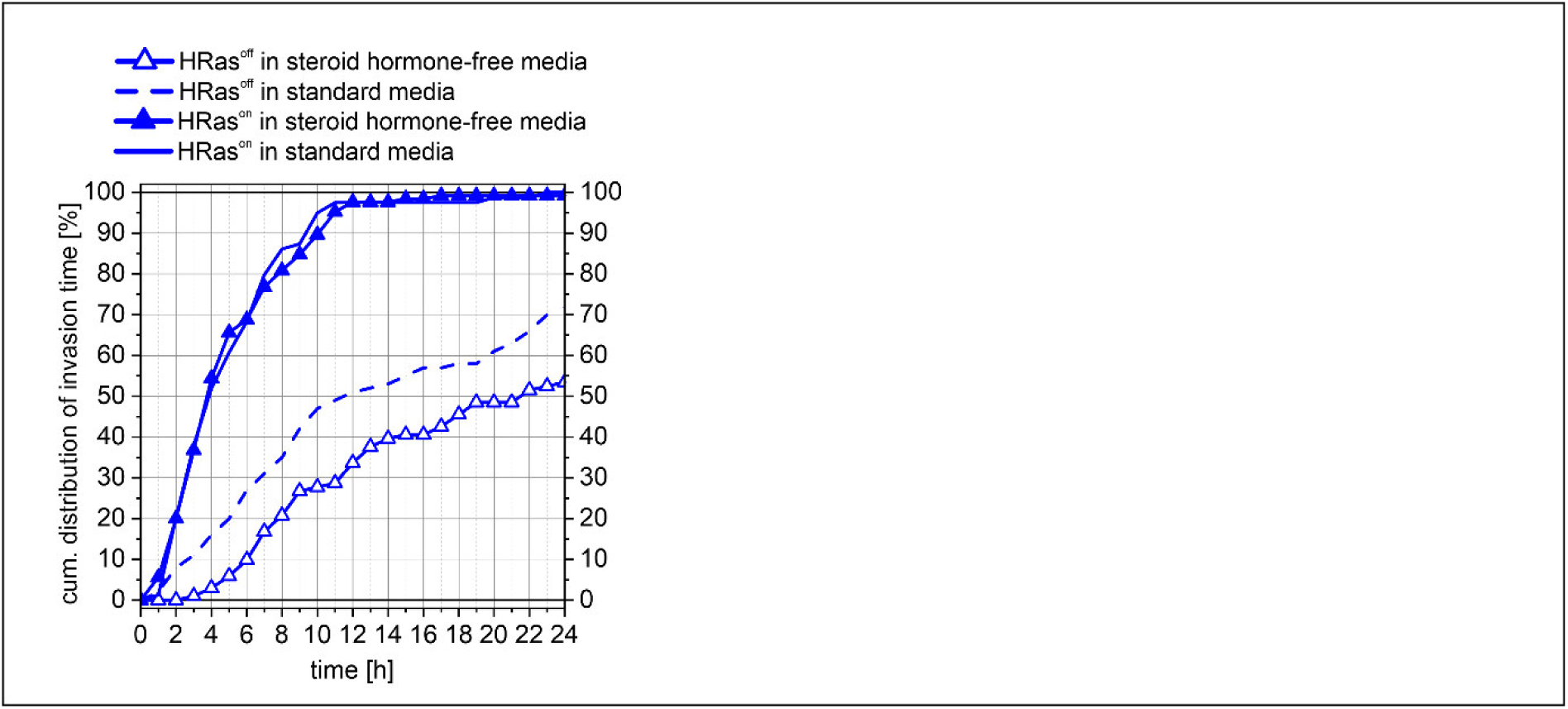
Steroid hormone-free media attenuates OHT-independent invasiveness of HRas^off^ spheroids. Cumulative distribution of BM transmigration time on stiff 16 kPa substrates (n ≥ 44 spheroids from ≥ 3 independent experiments). Data is shown here for comparison and corresponds to data in Fig 2D (standard media) and to data in Fig 6I (steroid-hormone-free media).

**Supplementary Figure S2:**
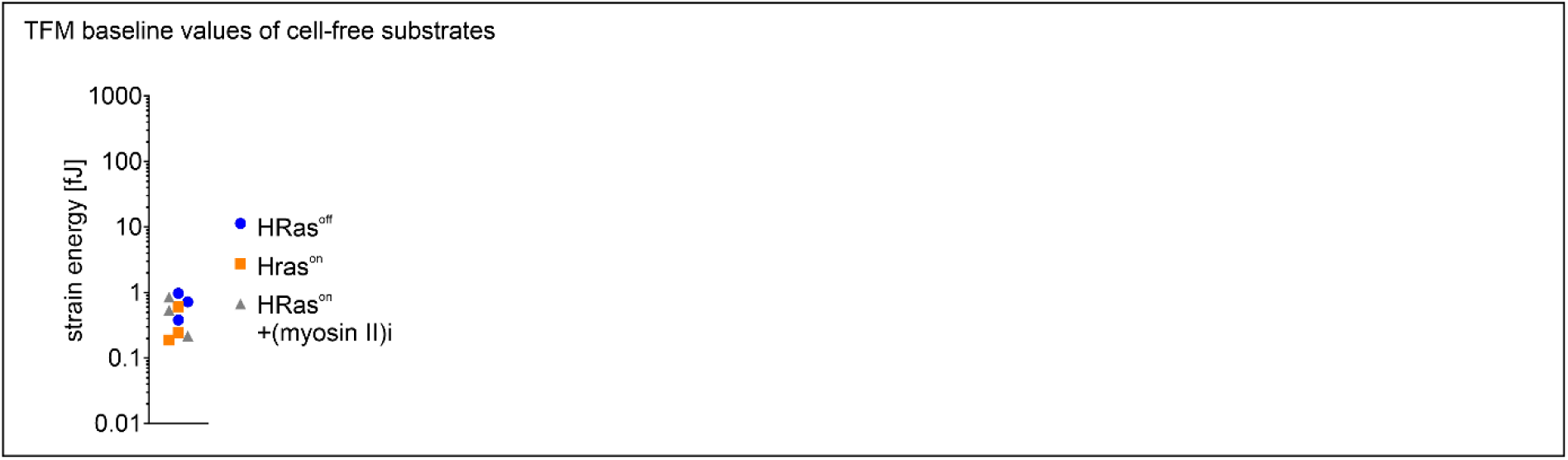
Substrate specific strain energy detected by traction force microscopy. Baseline of SE measured at cell free positions. For each condition, SE values were recorded at 5, 10, and 20 hours from at least 2 spheroid-free control positions per experiment and averaged to obtain experiment-specific baseline values. These mean baseline values are shown here (from 3 independent experiments per condition), indicated according to spheroid treatment.

**Supplementary Figure S3:**
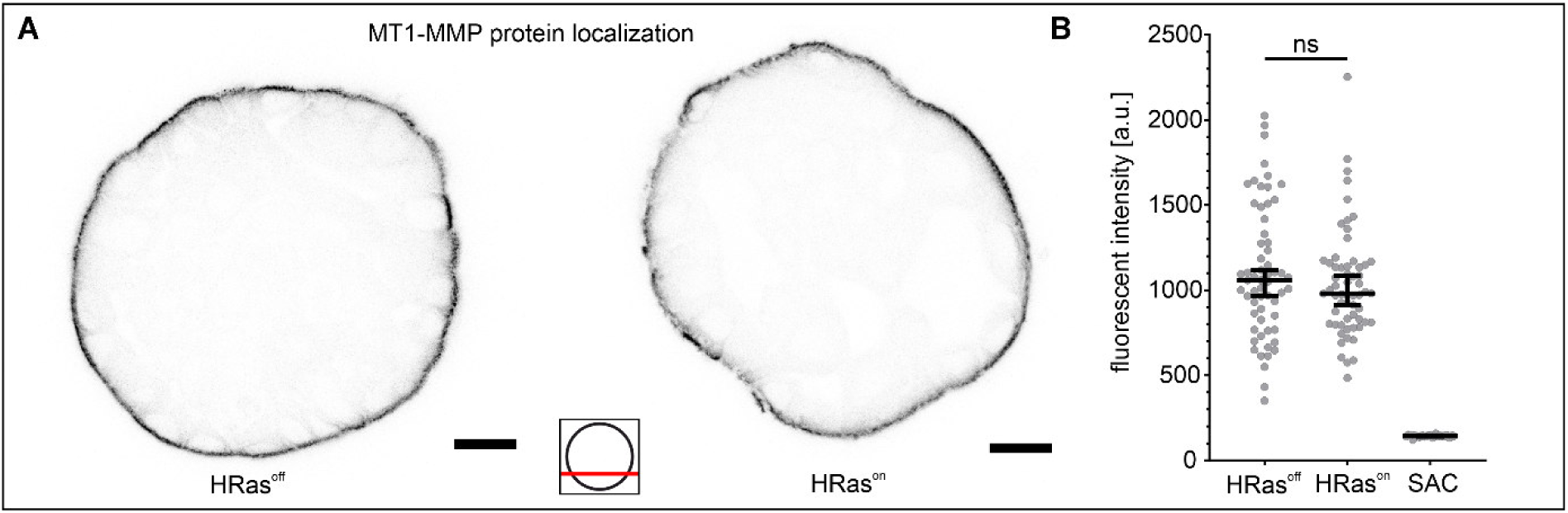
MMP localization and cellular content in HRas^on^ breast spheroids. (A) Representative images of MT1-MMP (inversed grey scale) staining of MCF10A/HRas spheroids incubated with 1 µM OHT or EtOH (HRas^off^ control) for 16 hours, and the secondary antibody control (SAC) to measure unspecific background signals. (B) Quantification of fluorescence intensities of MT1-MMP staining (n = 60 and n = 30 for SAC from three and two individual staining experiments, respectively). Scatter plot includes median and 95% CI. Mann-Whitney-U-test was performed for the data (n.s.: p > 0.05). Scale bars: 20 µm. Position of focal plane used for imaging and analyses is indicated by red bar.

**Supplementary Figure S4:**
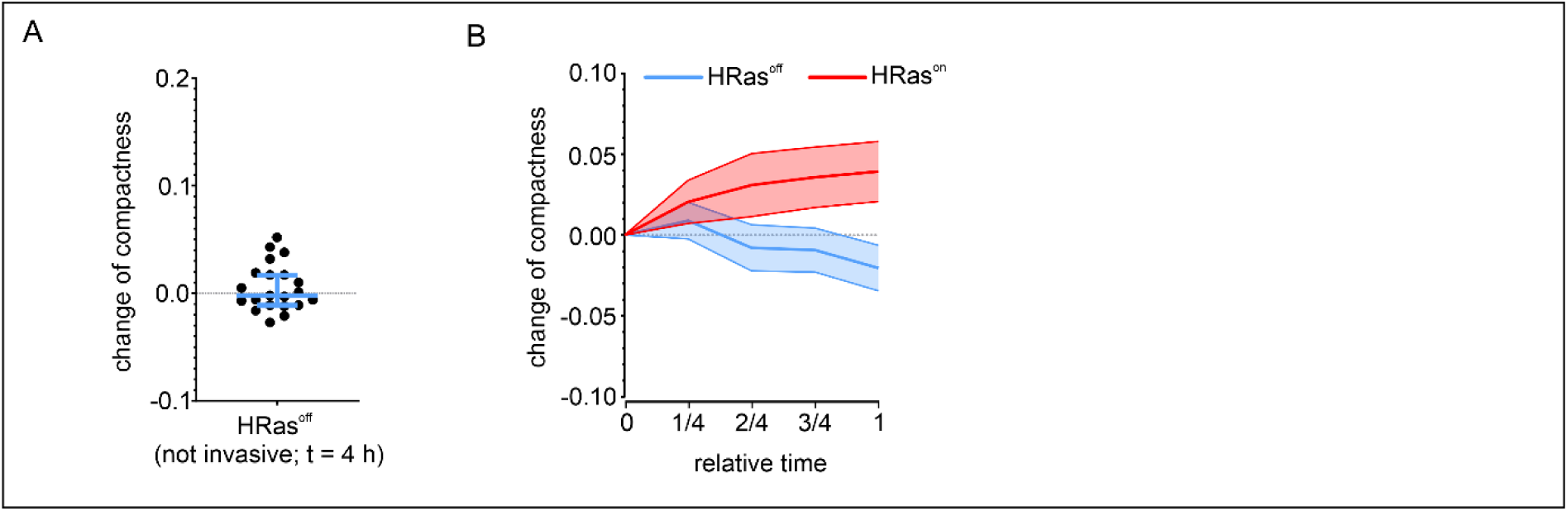
Comparative spheroid compactness depending on HRas activation. (A) Marginal changes of compactness for non-invasive HRas^off^ at 4 hours (= mean invasion onset of HRas^on^ group) control spheroids (n = 21 spheroids from 3 independent experiments). Shown with median and 95% CI. (B) Relative time curves show the change of compactness. To ease comparison data points were normalized: For non-invasive HRas^off^ spheroids, the overall measurements points at 3, 6, 9 and 12 hours were transformed to quarters 1/4, 2/4, 3/4 and 1, respectively. Each value was subtracted from the starting compactness (t = 0 hours). For invasive HRas^on^ spheroids the period until individual invasion onsets were quartered and subtracted from the respective starting compactness values (0 hours). Line represents the mean with 95% CI error bands (n ≥ 15 spheroids from ≥ 3 independent experiments).

**Supplementary Figure S5:**
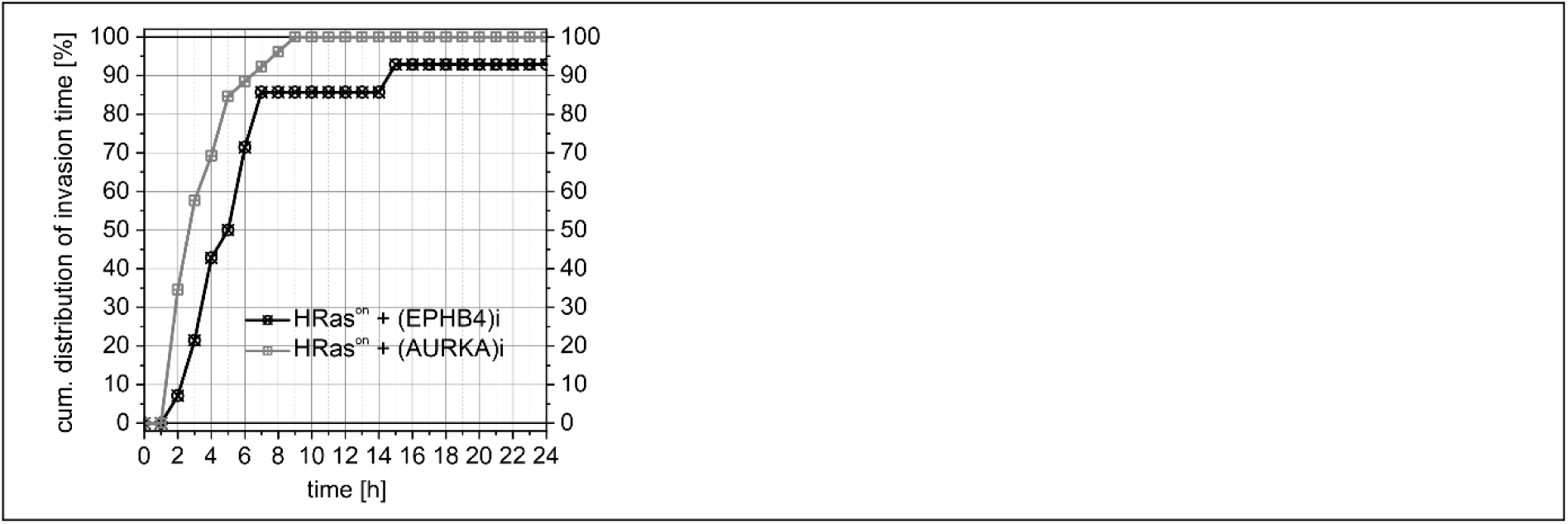
HRas-driven cell invasion is not dependent on EPHB4 or AURKA signaling. Cumulative distribution of BM transmigration time dependent on HRas induction and EPHB4-kinase and AURKA-kinase inhibition. Both experiments were done once. Higher inhibitor concentration showed cell toxicity.

**Supplementary Figure S6:**
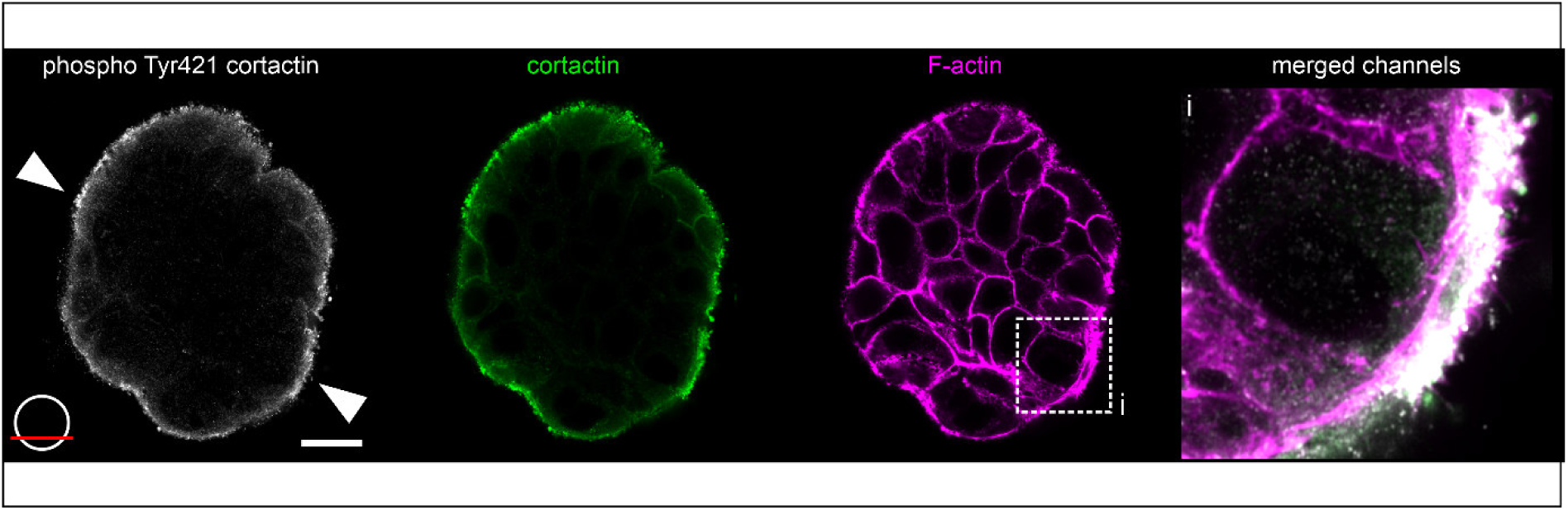
Co-localization of pTyr421 cortactin with reinforced cortical actin structures. Representative micrographs of fixed and immunostained HRas^on^ spheroid on 16 kPa substrate after 4 hours OHT treatment. Congruent pTyr42-cortactin (grey) and cortactin (green) signal distribution, accumulated at sites of thickened, reinforced actin (magenta), see white arrowheads. The zoom-in illustrates locally restricted phospho-cortactin-actin accumulation. Scale bar: 20 µm. Cartoon: Indicates The position of the optical image plane (red bar).

**Supplementary Figure S7:**
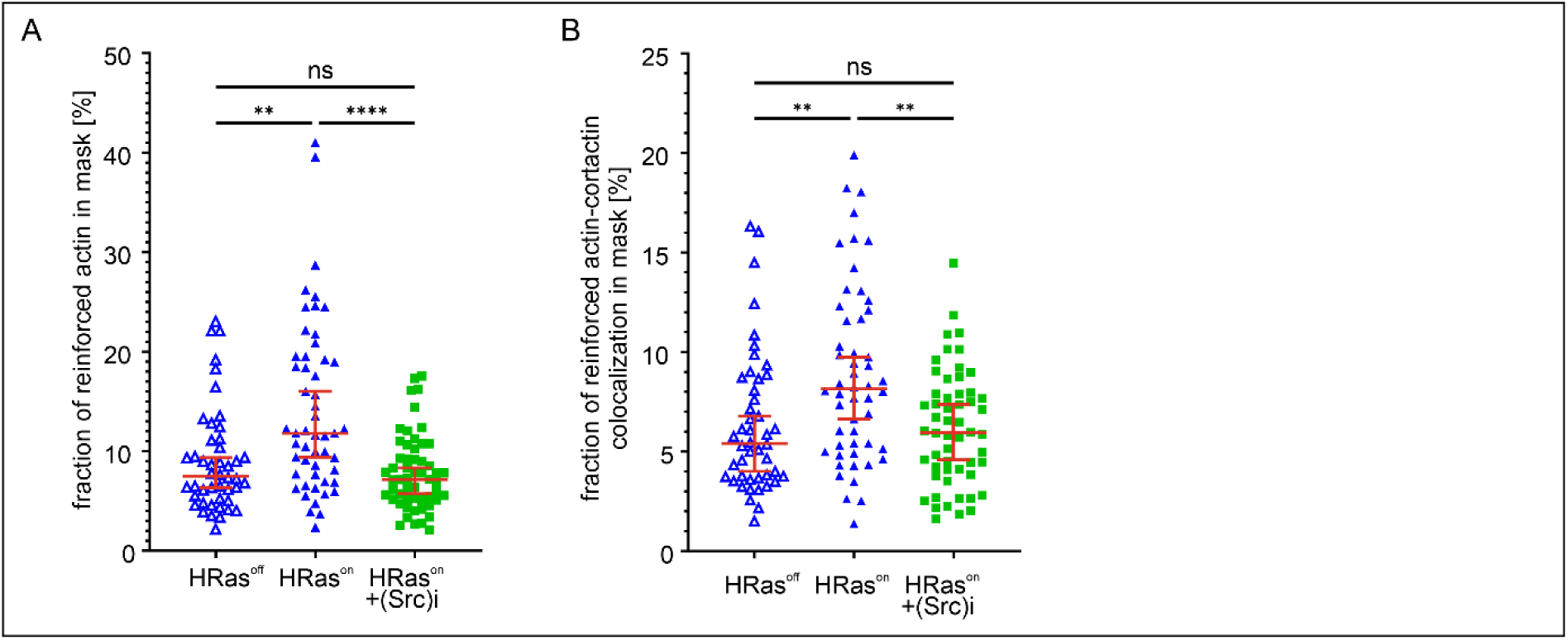
HRas activation increases actin reinforcement and cortactin co-localization. Coverage of reinforced actin patches [%] in (A) and additionally co-localized cortactin [%] in (B) of individual spheroids was plotted with median and 95% CI (n ≥ 46 for each sample condition, from 3 independent experiments). Kruskal-Wallis test with Dunn’s multiple comparison test was performed for the data (n.s.: p > 0.05; *: p ≤ 0.05; **: p ≤ 0.01; ***: p ≤ 0.001; ****: p ≤ 0.0001).

**Supplementary Figure S8:**
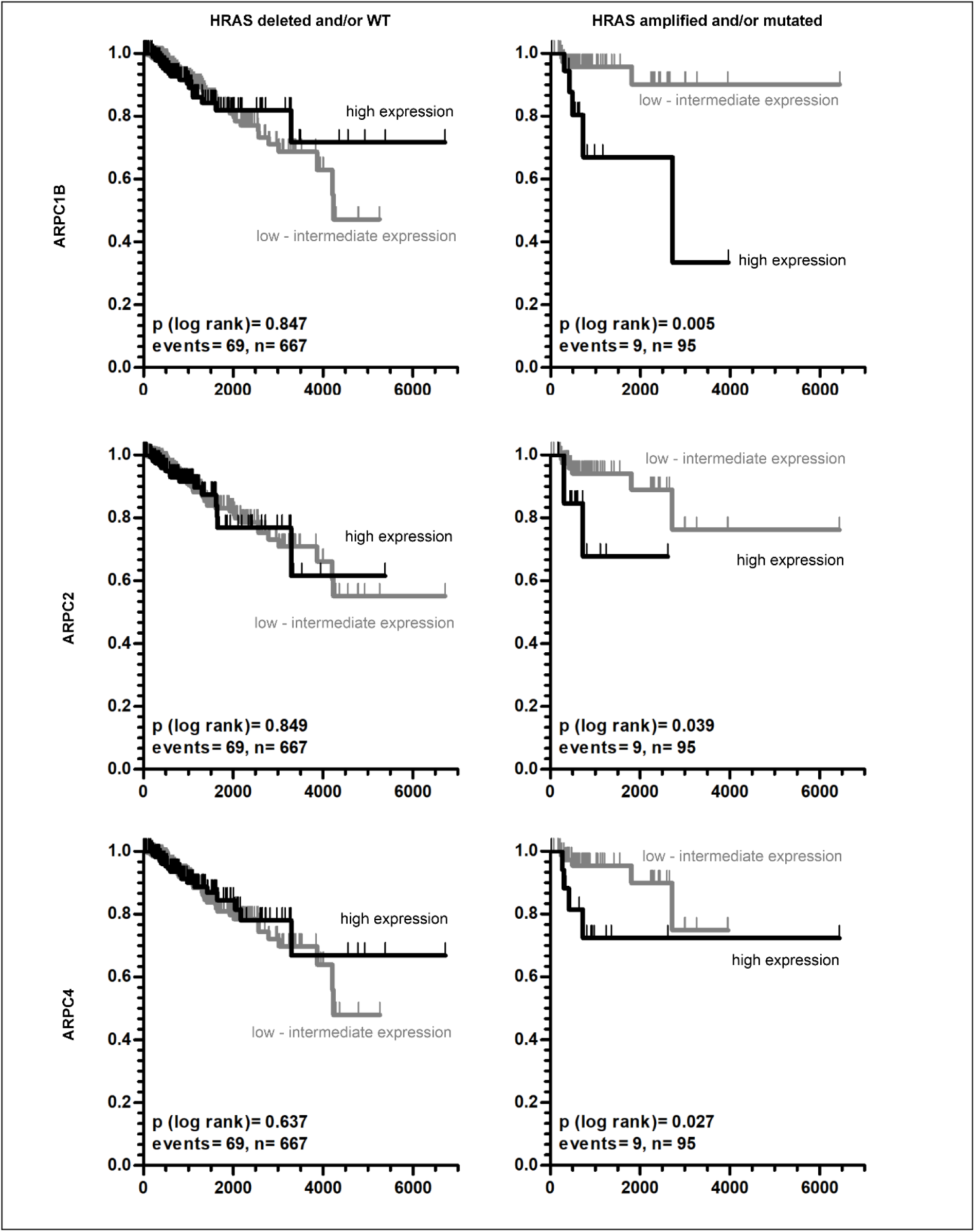
High ARP2/3 subunit and HRas gene expression predict poor survival of breast cancer patients. Univariate Kaplan-Meier curves show the recurrence-free survival (RFS) of breast cancer patients stratified by HRAS status (deletion or WT, left row; amplification or mutation, right row) RFS is shown depending on actin-related protein (ARP) subunits (HGNC group ID: 39) gene expression states (low vs. high).

## Supplemental Tables

**Supplementary table 1:**
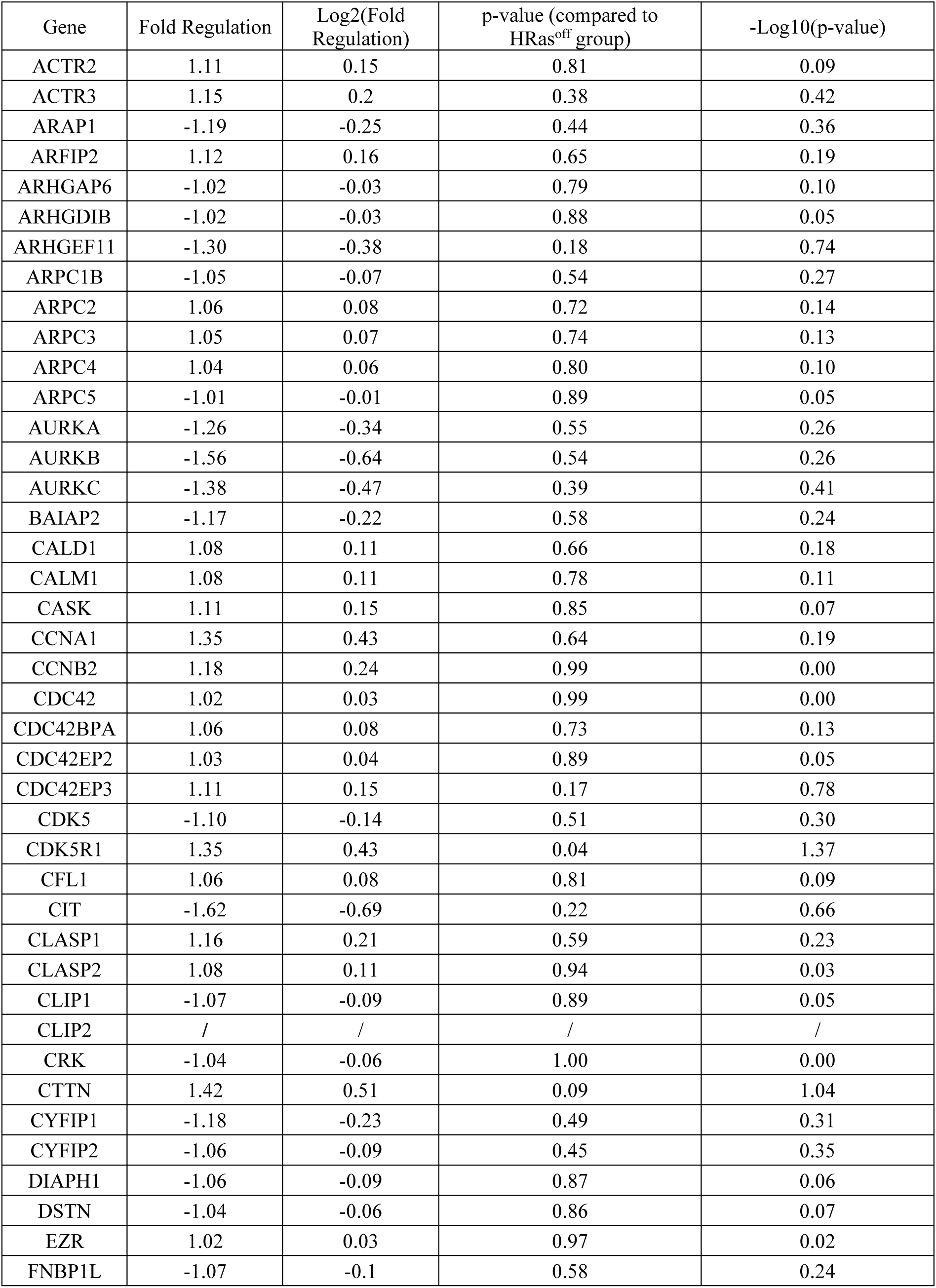
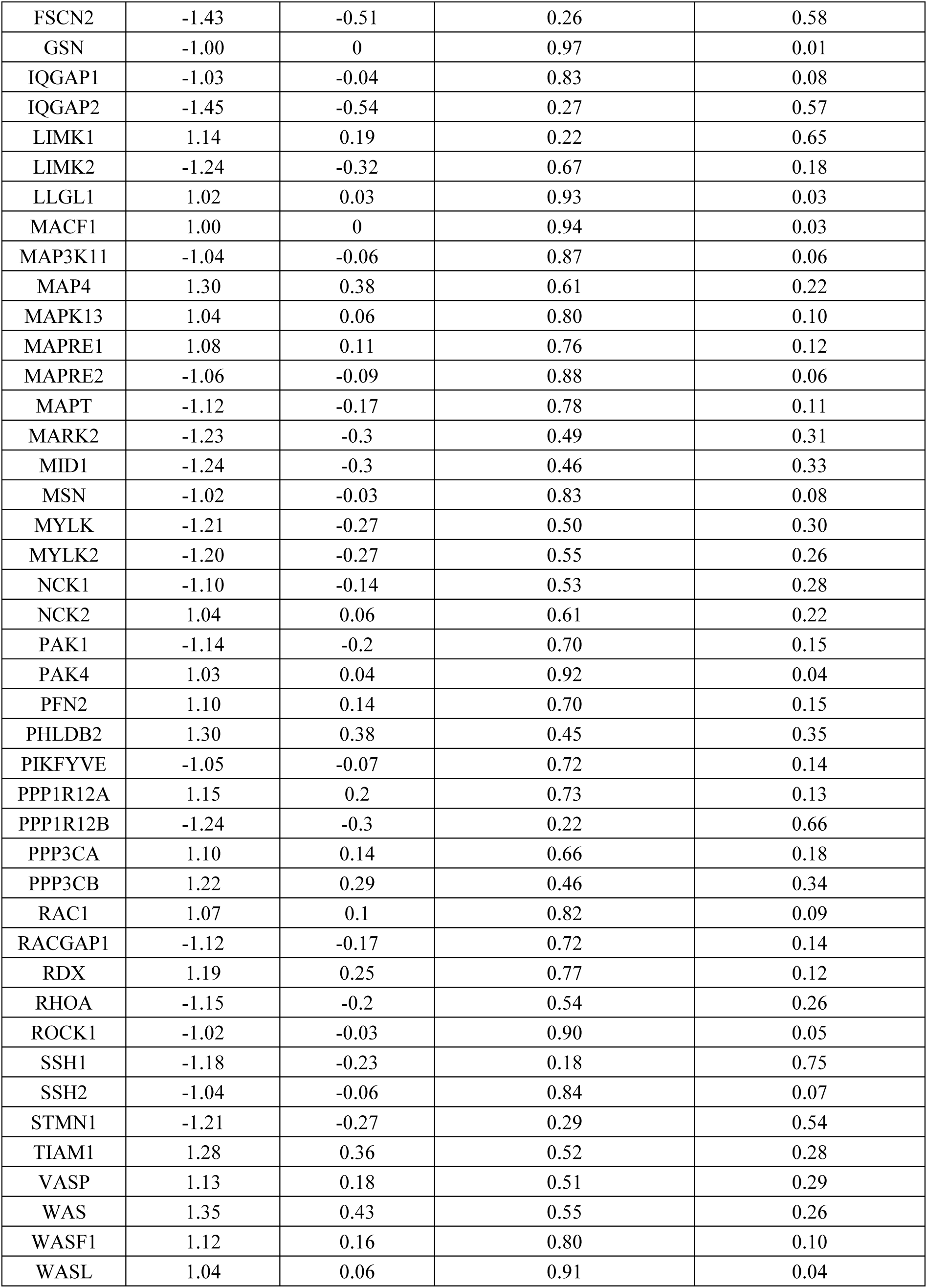
Raw data of mRNA expression values of 84 genes related to cytoskeletal regulation measured using a qRT-PCR array (QIAGEN, PAHS-088Z). Gene expression was analyzed in HRas^on^ spheroids after 4.5 hours of OHT treatment compared to HRas^off^ spheroids, both on tumor stiffness (16 kPa).

**Supplementary table 2:** Full data set of kinase profiling analysis of phospho-tyrosine-kinase (RTK) and serine-threonine-kinase (STK) activation depending on HRas activation retrieved from kinase-substrate microarrays. Kinase rank: decreasing p-values of change in kinase activity. Mean kinase statistic: color coded up (1)- and down (-1)-regulation. Kinase regulation in HRss^on^ spheroids (after 1 hour of OHT) was compared with HRas^off^ controls, all cultivated on 16 kPa substrates. PamChip® peptide tyrosine kinase (PTK) microarray system or the PamChip® Ser/Thr Kinase assay (STK).

| PTK |  |  |  | STK |  |  |  |
| --- | --- | --- | --- | --- | --- | --- | --- |
| Kinase Uniprot ID | Kinase Name | Kinase rank/Median Final score | Mean Kinase Statistic | Kinase Uniprot ID | Kinase Name | Kinase rank/Median Final score | Mean Kinase Statistic |
| P54760 | EphB4 | 2.27868 | 1.57606 | P31751 | Akt2/PKB[beta] | 2.01484 | 0.48170 |
| Q06187 | BTK | 1.83851 | 0.84904 | Q13464 | ROCK1 | 1.98851 | 0.74470 |
| P29317 | EphA2 | 1.79134 | 0.78815 | P68400 | CK2[alpha]1 | 1.91966 | 0.72202 |
| P07949 | Ret | 1.71652 | 0.91063 | Q9UBS0 | p70S6K[beta] | 1.86851 | 0.47515 |
| P07333 | FmS/CSFR | 1.68445 | 0.80322 | P31749 | Akt1/PKB[alpha] | 1.71381 | 0.45414 |
| P42681 | TXK | 1.56786 | 0.71669 | O14965 | AurA/Aur2 | 1.57551 | 0.67705 |
| P10721 | Kit | 1.51902 | 0.71311 | O75582 | MSK1 (RPS6KA5) | 1.57355 | 0.52124 |
| Q06418 | Tyro3/Sky | 1.50048 | 0.76366 | P49674 | CK1[epsilon] | 1.52719 | 0.57413 |
| Q08881 | ITK | 1.46223 | 0.73304 | Q15131 | CDK10 | 1.30515 | 0.45603 |
| P16591 | Fer | 1.44254 | 0.65002 | P51812 | RSK2 | 1.16236 | 0.38970 |
| P12931 | Src | 1.30047 | 0.71359 | Q15349 | RSK1/p90RSK | 1.16236 | 0.45373 |
| P54753 | EphB3 | 1.28231 | 1.01164 | P10398 | ARAF | 1.08933 | -0.50885 |
| P07332 | Fes | 1.27638 | 0.65748 | P15056 | BRAF | 1.06932 | -0.32671 |
| P08581 | Met | 1.02885 | 0.56433 | Q96GD4 | AurB/Aur1 | 1.06060 | 0.48304 |
|  |  |  |  | P51817 | PRKX | 1.05329 | 0.38842 |
|  |  |  |  | Q16566 | CaMK4 | 1.01226 | 0.42324 |

